# Molecular assembly of measles and Nipah virus: specific lipid binding drives conformational change and matrix polymerization

**DOI:** 10.1101/2021.10.11.463969

**Authors:** Michael J. Norris, Monica L. Husby, William B. Kiosses, Jieyun Yin, Linda J. Rennick, Anja Heiner, Stephanie Harkins, Rudramani Pokhrel, Sharon L. Schendel, Kathryn M. Hastie, Sara Landeras-Bueno, Zhe Li Salie, Benhur Lee, Prem P. Chapagain, Andrea Maisner, W Paul Duprex, Robert V. Stahelin, Erica Ollmann Saphire

## Abstract

Measles virus, Nipah virus, and multiple other paramyxoviruses cause disease outbreaks in humans and animals worldwide. The paramyxovirus matrix (M) protein mediates virion assembly and budding from host cell membranes. M is thus a key target for antivirals, but few high-resolution structures of paramyxovirus M are available, and we lack the clear understanding of how viral M proteins interact with membrane lipids to mediate viral assembly and egress needed to guide antiviral design. Here, we reveal that M proteins associate with phosphatidylserine and phosphatidylinositol-4,5-bisphosphate (PI(4,5)P_2_) at the plasma membrane. Using X-ray crystallography, electron microscopy, and molecular dynamics we demonstrate that PI(4,5)P_2_ binding induces conformational and electrostatic changes in the M protein surface that trigger membrane deformation, matrix layer polymerization, and virion assembly.

## Introduction

Paramyxoviruses are enveloped, negative-sense RNA viruses that are among the most infectious and pathogenic viruses known (Lamb and Parks, 2013; Rima et al., 2019). Most paramyxoviruses spread through respiratory droplets. Measles virus (MeV) still infects >7 million people and causes >100,000 deaths annually (Dabbagh et al., 2018; Portnoy et al., 2019). Multiple other paramyxoviruses cause devastating annual disease in humans, including Nipah virus (NiV) with up to 90% lethality (Amaya and Broder, 2020), and Parainfluenza virus III (PIV-III), a leading cause of childhood hospitalization (Abedi et al., 2016). Avian and ruminant paramyxoviruses threaten global food supplies and cause substantial annual economic losses (Brown and Bevins, 2017; Jones et al., 2016). Novel paramyxoviruses, to which humans are immunologically naive, may yet emerge with significant pandemic potential (Thibault et al., 2017). Despite the number and breadth of these threats, no specific therapies are yet available against any paramyxovirus.

Paramyxoviruses bud from the plasma membranes (PMs) of infected cells. Viral matrix (M) proteins coordinate budding of new virions by marshaling other viral structural components, including surface glycoproteins and viral replication complexes at assembly sites along the host PM. Expression of M alone can drive budding of virus-like particles (VLPs) of most paramyxoviruses (Harrison et al., 2010), while loss or mutation of M severely impairs viral replication (Cathomen et al., 1998; Dietzel et al., 2015; Ringel et al., 2020).

During infection, M proteins form a paracrystalline lattice at discrete assembly sites underlying infected cell PMs (Battisti et al., 2012; Ke et al., 2018a). These M lattices bridge viral glycoproteins and the internal ribonucleocapsid (RNP) complex containing the RNA genome to form the virion particle (Battisti et al., 2012; Harrison et al., 2010; Iwasaki et al., 2009; Ke et al., 2018a; Ray et al., 2016; Tahara et al., 2007). However, the nature of M protein binding to membranes, why virus assembly happens largely at the PM, what triggers viral matrix polymerization, and whether interaction of M with lipid membranes alone is sufficient to form outward protrusions in the PM, remain unclear.

Here, we show for the first time that phosphatidylserine (PS), and the phosphatidylinositol PI(4,5)P_2_ mediate MeV- and NiV-M protein interactions with host membranes. We describe the first high-resolution crystal structures for MeV- and NiV-M proteins, demonstrate that dimerization is critical for virion formation and illuminate structural rearrangements in M that occur upon binding of PI(4,5)P_2_ with a crystal structure of NiV-M protein bound to a soluble form of PI(4,5)P_2_. We further show that these lipid-induced structural rearrangements alter the M dimer shape and electrostatics, to promote M protein lattice polymerization and membrane curvature that together form the virion. The M-lipid complex structure also reveals new 3D templates for design of agents to inhibit paramyxovirus assembly.

## Results

### Electrostatic interactions govern NiV-M PM localization but not MeV-M

Paramyxovirus M proteins are known to localize at the PM inner leaflet during viral infection (reviewed in El Najjar et al., 2014), but the mechanism by which this localization occurs remains poorly characterized. Electrostatic interactions between the positively charged surface of M proteins and the negatively charged PM inner leaflet are hypothesized to facilitate membrane localization (Battisti et al., 2012; Liu et al., 2018). However, earlier studies indicated that shielding electrostatic interactions with high salt does not inhibit membrane binding of some paramyxovirus M proteins, suggesting that membrane localization may instead involve interaction with specific lipids (Faaberg and Peeples, 1988; Riedl et al., 2002; Stricker et al., 1994). To better understand the mechanism and specificity of M localization at the PM, we tested the impact of PM charge neutralization on M protein localization by treating cells with sphingosine, a positively charged lipid molecule that incorporates into the PM inner leaflet. The presence of sphingosine neutralizes the overall negative surface charge of the inner leaflet, and in turn substantially reduces nonspecific electrostatic interactions at the PM, but does not inhibit specific charge-based interactions (Yeung et al., 2008). Here we found that sphingosine does not affect PM localization of MeV-M (Figure 1A). This result is similar to that for eVP40 (Johnson et al., 2016), a control protein that interacts with specific lipid head groups (Figure S1A), suggesting that MeV-M PM localization involves interactions with specific lipids rather than nonspecific, negative charge-dependent membrane association. Meanwhile, in the presence of sphingosine NiV-M PM localization is reduced (Figure 1B). This reduction is similar to that seen for the control protein KRϕ-RFP (Yeung et al., 2006) that localizes to the PM via nonspecific electrostatic interactions (Yeung et al., 2006) (Figure S1B). Together, these results indicate that MeV- and NiV-M traffic to the PM largely through interactions with specific lipids and electrostatic associations, respectively.

**Figure 1.**
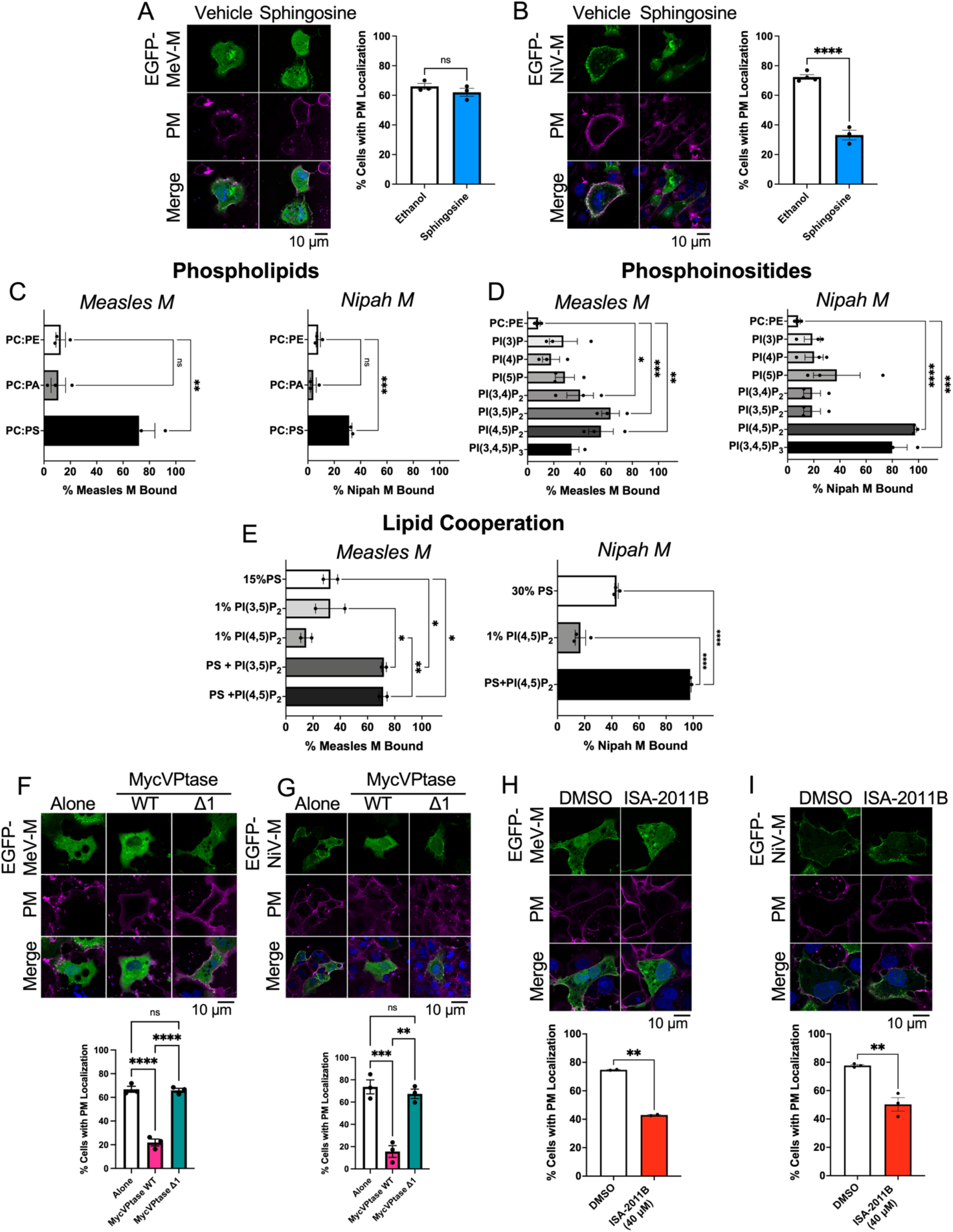
MeV-M and NiV-M preferentially interact with phosphatidylserine and PI(4,5)P_2_. (A & B; *Left*) COS-7 cells expressing the indicated GFP-fused protein for 24 hours, and treated with the membrane permeable base, sphingosine (37.5 µM) or ethanol vehicle (1:2000 v/v), for 1 hr at 37°C before staining and CLSM. Representative CLSM images are shown. (A & B; *Right*) Quantification of PM localization. Cells were counted and binned based on high or no fluorescent signal at the PM and the percentage of cells with PM localization was calculated (N≥45 cells/replicate). Liposome sedimentation assays and western blotting of MeV- or NiV-M binding to LUVs containing (C) phospholipids or (D) phosphatidylinositols. (E) MeV-M and NiV-M affinity for liposomes containing different lipid combinations. The percent bound M was determined by densitometry of bands for the supernatant and pellet fractions (Figure S1C-E) (N≥5) (F & G) *Top:* Representative CLSM of COS-7 cells expressing EGFP-MeV-M (F) or EGFP-NiV-M (G) alone (left panels) or with MycVPtase-WT (middle panels) or MycVPtase-Δ1 (right panels). (H & I) *Top:* ISA-2011B (40 µM or DMSO vehicle 1:1000 v/v) treatment of COS-7 cells expressing EGFP-MeV-M (H) or EGFP-NiV-M (I) 8 hours post-transfection for 24 hrs at 37°C. (F-I; *Bottom*) PM localization of MeV- and NiV-M quantified as per right panels of A & B. For all CLSM images (A & B; F-I) cells were stained 24 hr post-transfection with WGA AlexaFluor647 and Hoechst (PM and DNA stain, respectively) before imaging. Unless otherwise stated, Scale bar=10 µm. Values are the mean ± SEM of 3 independent experiments. A one-way ANOVA with Tukey’s *post hoc* test for group comparisons was performed or a Student’s t-test to compare treatment to the vehicle group for each protein was performed. ns P > 0.05, *P ≤ 0.05, **P ≤ 0.01, ***P ≤ 0.001, ****P ≤ 0.0001. PC: phosphatidylcholine; PE: phosphatidylethanolamine; PA: phosphatidic acid; PS: phosphatidylserine; PIP: phosphatidylinositol-phosphate; MycVPtase: Myc-5–phosphatase; WGA: wheat germ agglutinin; PM: plasma membrane; CLSM: confocal laser scanning microscopy.

### Measles and Nipah M proteins selectively bind phosphatidylserine and PI(4,5)P_2_ in membranes

Although paramyxovirus M proteins intrinsically bind cellular membranes during assembly, whether particular lipid head groups anchor M to the inner leaflet is unclear (Riedl et al., 2002; Shnyrova et al., 2007; Stricker et al., 1994; Subhashri and Shaila, 2007). To understand how MeV- and NiV-M anchor to membranes, we used liposome sedimentation assays to identify specific lipids critical for membrane binding. No significant binding of MeV- or NiV-M to large unilamellar vesicles (LUVs) comprising phosphatidylcholine (PC) and phosphatidylethanolamine (PE) or negatively charged phosphatidic acid (PA) was observed (Figure 1C; S1C). MeV- and NiV-M did, however, associate with LUVs carrying anionic phosphatidylserine (PS) (Figure 1C; S1C).

Phosphoinositides (PIs) are a minor lipid species in cell membranes, yet play vital roles in membrane trafficking and cell signaling. There are seven PI species present in eukaryotic membranes (Hammond and Burke, 2020). MeV- and NiV-M each had only minimal binding to LUVs carrying mono-PIs (PI3P, PI4P, and PI5P; Figure 1D & S1D). MeV-M exhibited significant binding to LUVs containing the bi-PIs PI(3,5)P_2_ or PI(4,5)P_2_ and NiV-M interacted with LUVs containing PI(4,5)P_2_, or the tri-PI PI(3,4,5)P_3_ (Figure 1D; S1D).

### M proteins cooperatively bind PI(4,5)P_2_ and PS

Peripheral proteins can interact with multiple lipids in the PM to increase protein-membrane affinity (Cho and Stahelin, 2005). Indeed, we find that inclusion of PS in LUVs containing PI(4,5)P_2_ enhances MeV- and NiV-M binding by ∼2.5 and 5 fold, respectively, compared to LUVs with PI(4,5)P_2_ alone (Figure 1E; S1E). PS and PI(4,5)P_2_ are the predominant anionic lipids in the PM (Yang et al., 2018), suggesting that synergistic binding with the viral matrix could facilitate PM anchoring and subsequent M-driven viral budding.

### PI(4,5)P_2_ is essential for measles and Nipah M membrane interaction in live cells

We then investigated if MeV- or NiV-M PI(4,5)P_2_ association detected in liposome sedimentation assays was functionally significant in a cellular system in which PI(4,5)P_2_ primarily localizes to the PM inner leaflet (Burke, 2018). We co-expressed EGFP-MeV-M or EGFP-NiV-M with a phosphoinositide 5-phosphatase (MycVPtase) that cleaves the phosphate from the D5 position of PI(4,5)P_2_ to reduce PI(4,5)P_2_ levels in the PM, and determined M protein localization. Inactive MycVPtase (MycVPtase-Δ1) and proteins that interact with PI(4,5)P_2_ (PLC***δ***-PH & VP40) or PS (LactC2) were used as controls (Figure S2A-C). We found that MycVPtase-dependent PI(4,5)P_2_ depletion significantly reduces both EGFP-MeV-M and EGFP-NiV-M PM localization (Figure 1F & G). Similarly, the PI-5-kinase-α (PIP5kα) inhibitor ISA-2011B, which inhibits production of PI(4,5)P_2_ from its PI(4)P precursor, also reduced EGFP-MeV-M and EGFP-NiV-M PM localization (Figure 1H & I).

Like PI(4,5)P_2_, PI(3,4,5)P_3_, which is uniquely bound by NiV-M, is also found in the PM inner leaflet (Burke, 2018). The phosphatidylinositol 3-kinase (PI3K) inhibitor wortmannin reduces PI(3,4,5)P_3_ levels in the PM (Powis et al., 1994). Our results suggest that NiV-M PM localization is independent of PI(3,4,5)P_3_ as evidenced by the lack of effect by wortmannin (Figure S2D-F).

Conversely, PI(3,5)P_2_, preferred by MeV-M, is primarily found in multivesicular bodies (MVBs) (Burke, 2018). The phosphatidylinositol-3-phosphate 5-kinase (PIKfyve) inhibitor apilimod reduces cellular levels of PI(3,5)P_2_ (Sbrissa et al., 2018). Here apilimod-dependent reduction of PI(3,5)P_2_ in intracellular membranes decreased EGFP-MeV-M PM localization, which highlights a potential role for PI(3,5)P_2_ in MeV-M trafficking to the PM (Figure S2G-J). Together, these data suggest that PI(4,5)P_2_ is the primary driver of membrane interaction for both MeV- and NiV-M. Additionally, PI(3,5)P_2_ may enhance interaction of MeV-M with MVBs.

### PI(4,5)P2 drives conformational changes in paramyxovirus M dimers

Since PI(4,5)P_2_ was important for PM localization and anchoring of both MeV- and NiV-M, we next explored the nature of this lipid interaction at a molecular level. We determined the crystal structures of both M proteins alone and the crystal structure of NiV-M in complex with PI(4,5)P_2_. Consistent with other paramyxovirus M proteins (Battisti et al., 2012; Liu et al., 2018) both Apo forms of MeV- and NiV-M form head-to-tail dimers, stabilized by hydrophobic and electrostatic interactions, with buried surface of ∼2300 Å^2^ (Figure S3A & B; Table S1). The protomers have two structurally analogous domains joined by a flexible linker region. A twisted β-sandwich surrounded by several short α-helices forms the core of each domain. Both MeV- and NiV-M crystal structures exhibit a predominant positive surface charge, and a large, uniform surface-exposed basic patch in each carboxyl-terminal domain (CTD) (Figure S3C & D) Size-exclusion chromatography coupled to multiangle light scattering (SEC-MALS) analysis and single particle negative stain electron microscopy (nsEM) further confirm the dimeric nature of MeV- and NiV-M in solution (Figure S3E-J).

The M-lipid crystal structure (Figure 2A; Table S1) reveals that PI(4,5)P_2_ binding induces extensive conformational rearrangements (Figure S3K) that further stabilize the dimer, flatten the membrane-interacting surface from the concave shape of the Apo structure, and alters the surface electrostatics. Specifically, lipid binding expands the dimer interface by 33 residues compared to Apo-NiV-M, likely increasing matrix lattice stability. Expansion of the interprotomer interaction in the lipid-bound dimer occurs as α-helix 1 in the Apo-NiV-M dimer interface unwinds. This unwinding increases mobility of the C-terminal downstream 20 residues, which rearrange from their initial position behind their own protomer to thread forward and reach across to a hydrophobic pocket in the NTD of the other protomer (Figure 2B). Although the N-terminal 30 residues are disordered in the Apo-NiV-M structure, they can be modeled in the lipid-bound NiV-M structure and occupy the region behind the protomer which was previously occupied by the C-terminus in the Apo-NiV-M structure (Movie S1). Structural rearrangements induced by PI(4,5)P_2_ binding also increase the angle between adjacent protomers (Figure 2C). This transition from the bowl-shaped Apo to the flatter, plate-shaped surface of the bound conformation could drive localized negative membrane curvature to facilitate initial membrane deformation at budding sites.

**Figure 2.**
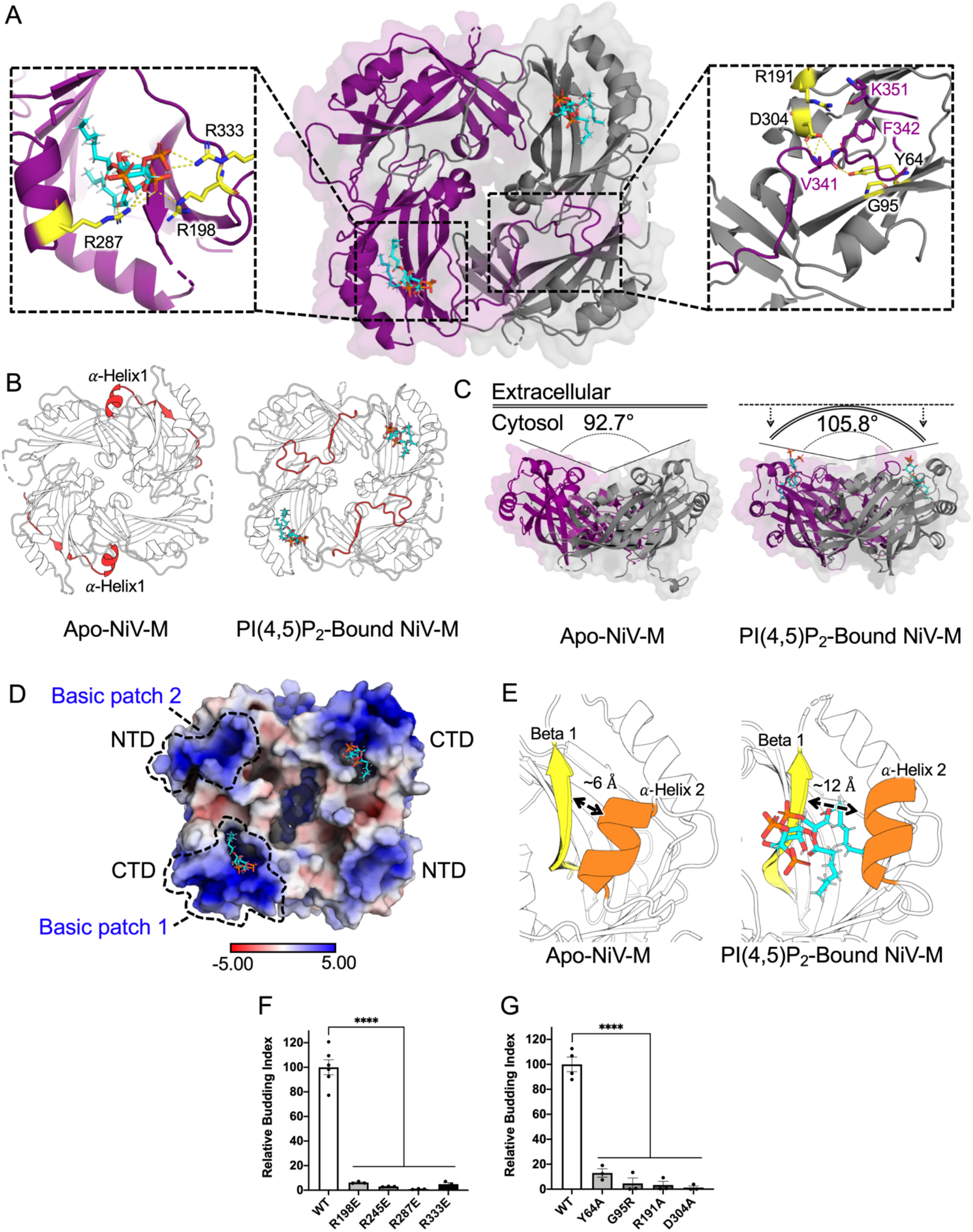
Binding of PI(4,5)P_2_ drives conformational changes in Nipah virus M protein. (A) Crystal structure of NiV-M dimer in complex with C8-PI(4,5)P_2_. NiV-M protomers are colored purple and grey and C8-PI(4,5)P_2_ molecules are colored cyan, orange, and red (for carbon, phosphorus, and oxygen atoms). *Left inset*: close-up view of PI(4,5)P_2_ binding site in the NiV-M CTD. *Right inset*: Conformational change in C-terminal residues (purple) extending across the hydrophobic pocket in the NTD of the adjacent protomer (grey). Main coordinating residues are highlighted yellow and shown as sticks. (B) Apo-NiV-M (left) and C8-PI(4,5)P_2_ bound NiV-M (right) showing the conformational change in the C- terminal 20 residues (red). (C) Side view of Apo-NiV-M (left) and C8-PI(4,5)P_2_-bound NiV-M (right). The angle between protomers increases as NiV-M transitions from a concave to flat surface after PI(4,5)P_2_ binding. (D) Electrostatic surface potential of C8-PI(4,5)P_2_-bound NiV-M (calculated by APBS) shown on a space-filling model with positively and negatively charged regions shown in blue and red, respectively. The CTD basic patch and newly formed basic patch in the NTD are outlined. (E) Close-up of PI(4,5)P_2_ binding pocket in Apo-NiV-M (left) and C8-PI(4,5)P_2_ bound NiV-M (right). The separation between α helix 2 and β sheet 1 increases by 6 Å in the conformational change required to accommodate lipid binding. (F & G) Comparison of VLP budding between WT and mutants targeting PI(4,5)P_2_- (F) or C-terminus-coordinating (G) residues of NiV-M. Normalized budding index was calculated from integrated intensities of the immunoblots (Figure S5B & C). Values represent mean ± SEM of three independent experiments. ****P ≤ 0.0001 by one-way ANOVA.

The conformational change also opens one basic patch and creates a second compared to the Apo form (compare Figure 2D and S3D). The first patch opens upon displacement of α-helix 2 from CTD β-sheet 1 to accommodate the negatively charged PI(4,5)P_2_ molecule (Figure 2E), which is coordinated by R198, R287, and R333 (Figure 2A; left inset). R333 is part of α-helix 1 in Apo-NiV-M, but is displaced by ∼7.6 Å in the bound form to coordinate PI(4,5)P_2_. Additionally, the PI(4,5)P_2_ acyl chains both pack against hydrophobic side chains in the CTD basic patch (Figure 2A; S3L). The second basic patch forms as the basic C-terminus drapes across the neighboring protomer NTD (compare Figure 2D and S3D). This second basic patch increases available surface area for interactions with negatively-charged lipid head groups, and can generate binding sites for membrane lipids like PS.

### Budding requires M protein dimerization

Previous results with related paramyxovirus M proteins suggested that dimerization is critical for virion formation (Battisti et al., 2012; Liu et al., 2018). To determine the role of M dimerization in MeV and NiV assembly, we introduced destabilizing mutations at the dimer interface of MeV- and NiV-M (Figure S3A & B). We confirmed the selected mutations disrupt the dimer interface using a bimolecular fluorescence complementation (BiFC) assay (Kerppola, 2006, 2008). Overall expression levels of each mutant were similar to wildtype (WT) M (Figure S4A & B). However, both confocal laser scanning microscopy (CLSM) (Figure S4A & B) and flow cytometry (Figure S4C & D) revealed mutant MeV- and NiV-M BiFC signals are diminished. To examine the role of M dimerization we quantified VLP production from mutant and WT M proteins. WT MeV- and NiV-M were detected both in total cell lysates and culture medium of mammalian cells, confirming that MeV- and NiV-M expression alone can drive VLP formation (Figure S4E & F). Meanwhile, VLP formation for MeV- and NiV-M mutants was substantially reduced relative to WT (Figure S4E & F). These results demonstrate that M dimerization is critical for virion formation.

### Requirement for PI(4,5)P_2_ binding and subsequent binding-induced conformational changes in M for budding

We next made charge-switching Arg to Glu mutations (R198D, R287D, and R333D) in the newly identified CTD basic pocket that receives PI(4,5)P_2_, and also at R245, which is disordered in the structure, but could participate in PI(4,5)P_2_ coordination as suggested by all-atom molecular dynamics simulations with PI(4,5)P_2_ (Figure S5A). In western blot analysis of VLP production in mammalian cells transfected with WT and mutant NiV-M, no NiV-M mutant promoted VLP production, indicating the importance of each Arg in the newly-formed PI(4,5)P_2_ pocket for NiV-M-mediated virus assembly and budding (Figure 2F; S5B).

To disrupt hydrogen bonding to the C-terminal mainchain during conformational rearrangement, we next mutated Y64, R191, D304 and the highly conserved G95. In the Apo-NiV-M structure, these residues are solvent-exposed and not readily involved in protein folding or M dimer formation. These mutants also lost all budding activity (Figure 2G; S5C), suggesting that interactions made by the NiV-M C-terminal 20 residues following conformational rearrangement upon PI(4,5)P_2_ binding are indeed a prerequisite for assembly and budding. In a similar panel of mutants for MeV-M, VLP formation decreased by up to 60% (Figure S5D-F), suggesting that similar molecular changes occur in MeV-M upon lipid binding.

### Modeling the plasma membrane association of NiV-M and MeV-M

To simulate membrane association between Apo-(Figure S3B) and lipid-bound (Figure 2A) conformations of NiV-M, we performed all-atom molecular dynamics (MD) simulations using realistic membrane bilayers containing POPC, POPS, POPE, palmitoylsphingomyelin (PSM), cholesterol, and PI(4,5)P_2_ at relevant ratios to model the PM in terms of charge, lipid packing, and hydrophobic core structure. Before the simulation, and after minimization and equilibration, both protein conformations were ∼10 Å below the PM lower leaflet and only weak interactions between proteins and membranes occurred (Figure 3A & B; 0 µs). For the Apo-NiV-M conformation, one M protomer interacted with the membrane for ∼0.040 µs before the second protomer began interacting. By ∼0.20 µs, many basic CTD residues in Apo-NiV-M interacted with anionic lipid headgroups, and by the end of the 0.5 µs simulation 35 lipids interacted with the dimer (Figure 3A; Movie S2). In contrast, the PI(4,5)P_2_-bound NiV-M conformation interacted rapidly with membrane, and by ∼0.06 µs most basic CTD residues in both protomers and several residues in the C-terminus that drapes across the adjacent protomer interacted with the membrane. By the end of the 1 µs simulation, 43 lipids interacted with the NiV-M dimer in the PI(4,5)P_2_-bound conformation (Figure 3B; Movie S2).

**Figure 3.**
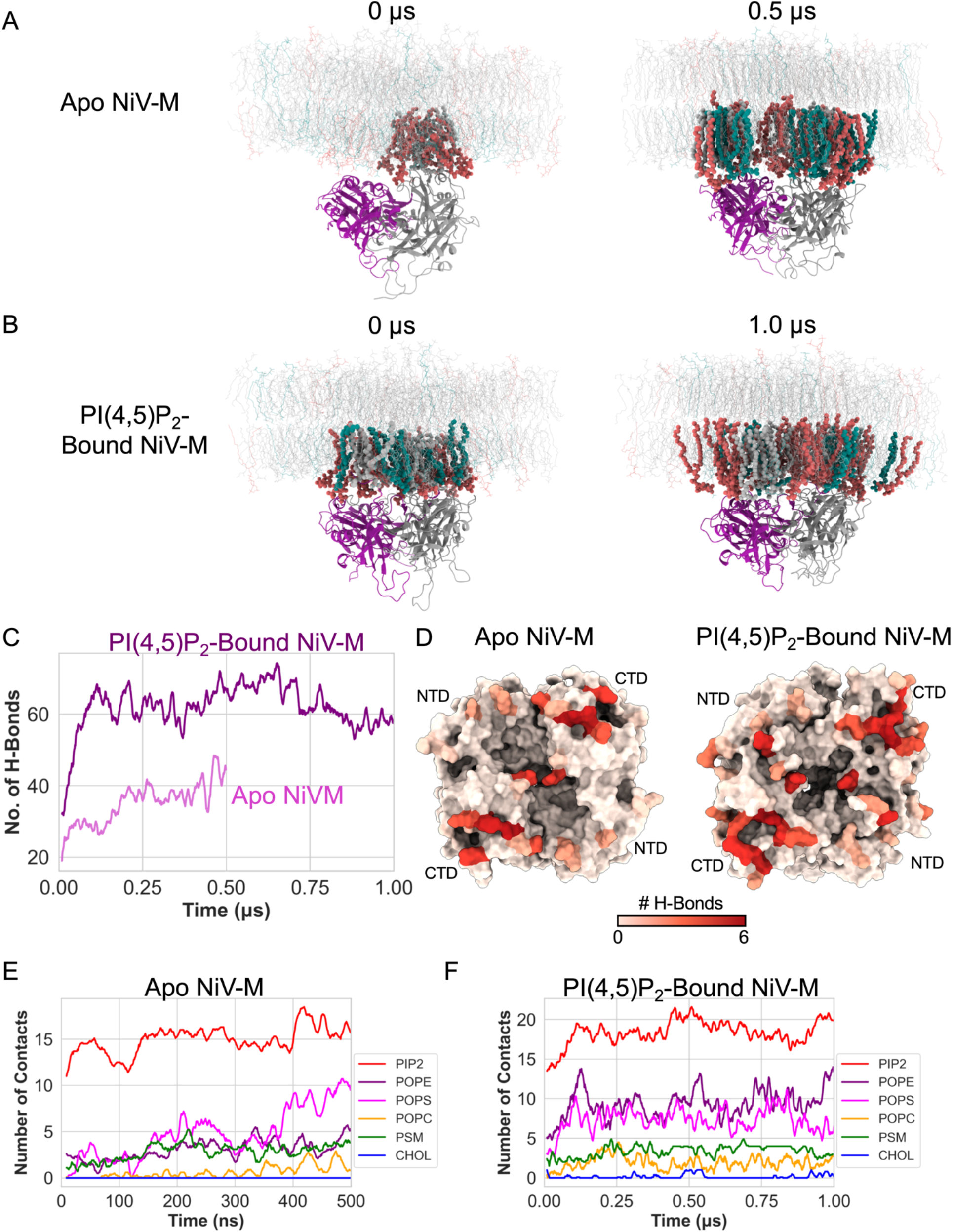
Molecular dynamics simulations of NiV-M interaction with the plasma membrane. Initial and final snapshots of NiV-M association with the PM in the (A) Apo and (B) PI(4,5)P_2_-bound conformation observed in the simulations. Left and right panels display representative initial and final configurations, respectively. Lipids interacting with protein are highlighted (PI(4,5)P_2_, red; PS, green; Cholesterol, POPC, POPE, and palmitoylsphingomyelin, grey). (C) Time evolution of hydrogen bond formation between membrane lipids and Apo (light purple) or PI(4,5)P_2_-bound (dark purple) NiV-M. (D) Surface model of Apo-NiV-M (left) and PI(4,5)P_2_-bound NiV-M (right). Residue colors correspond to the number of hydrogen bonds formed with the membrane within the last 200 ns of the simulation. (E & F) Time evolution of hydrogen bonds between various lipid types and the Apo-NiV-M (E) or PI(4,5)P_2_-bound conformation (F).

The PI(4,5)P_2-_bound-NiV-M had nearly 2-fold more hydrogen bonds with lipid headgroups than Apo-NiV-M during the MD simulation (Figure 3C). The rapidity of membrane interaction and increased number of membrane interactions further support that the conformational change in NiV-M observed in the PI(4,5)P_2_-bound NiV-M crystal structure (Figure 2) favors membrane association and protein stabilization at the PM lower leaflet. In the Apo-NiV-M conformation, hydrogen bonding with the PM involves almost exclusively CTD Arg residues (Figure 3D), whereas hydrogen bonding of the lipid-bound conformation also involves CTD Arg residues, including R198, R245, R287, and R333 (Figure 2F), as well as additional interactions with R244, K286, and R261. Extensive hydrogen bonding between membrane headgroups and both the C-terminal R348 that drapes across the NTD and the NTD R57 present in the second basic patch were also observed (Figure 3D).

We next calculated the number of lipid-protein contacts as a function of time (Figure 3E & F). Almost all lipid types bind to M proteins in both the Apo and bound conformations, but PS forms more contacts to PI(4,5)P_2_-bound NiV-M than the Apo conformation (Figure 3E & F). Further, PI(4,5)P_2_ (colored red in Figure 3A & B) makes substantially more interactions with basic residues than do other lipids. These results agree well with our experimental observations that PI(4,5)P_2_ and PS are important for PM localization and interaction (Figure 1). Using a homology model of bound-MeV-M generated from bound-NiV-M coordinates to perform the same MD simulations, lipid-bound MeV-M also forms significantly more hydrogen bonds with the PM and substantially more contacts with PI(4,5)P_2_ and POPS than the Apo-MeV-M conformation (Figure S6; Movie S3).

### Matrix proteins induce membrane deformation and curvature

As nascent virions form, the PM lipid bilayer curves outward (Welsch et al., 2007), but whether the matrix alone achieves this membrane curvature, or viral surface glycoproteins or cellular proteins are also involved is unclear (El Najjar et al., 2014; Harrison et al., 2010). To examine this membrane curvature we incubated MeV- or NiV-M with giant unilamellar vesicles (GUVs) lacking or containing PI(4,5)P_2_ and analyzed structural alterations in the lipid bilayer by CLSM. Both MeV- and NiV-M produced significant membrane deformation characterized by filamentous, spherical, and flat membrane protrusions in GUVs containing PI(4,5)P_2_ (Figure 4A & B). As the M proteins are outside the GUVs, all protrusions pointed inward to mimic negative curvature observed during virus budding. GUVs containing PS, but only trace amounts of PI(4,5)P_2_, remain spherical in the presence of MeV- or NiV-M, except at the highest M protein concentration tested (Figure 4A & B). Thus, at low protein densities, interactions between M proteins and PS alone do not appear to induce membrane deformation, whereas the conformational changes we observed upon interaction with PI(4,5)P_2_ (Figure 2) can induce membrane curvature.

**Figure 4.**
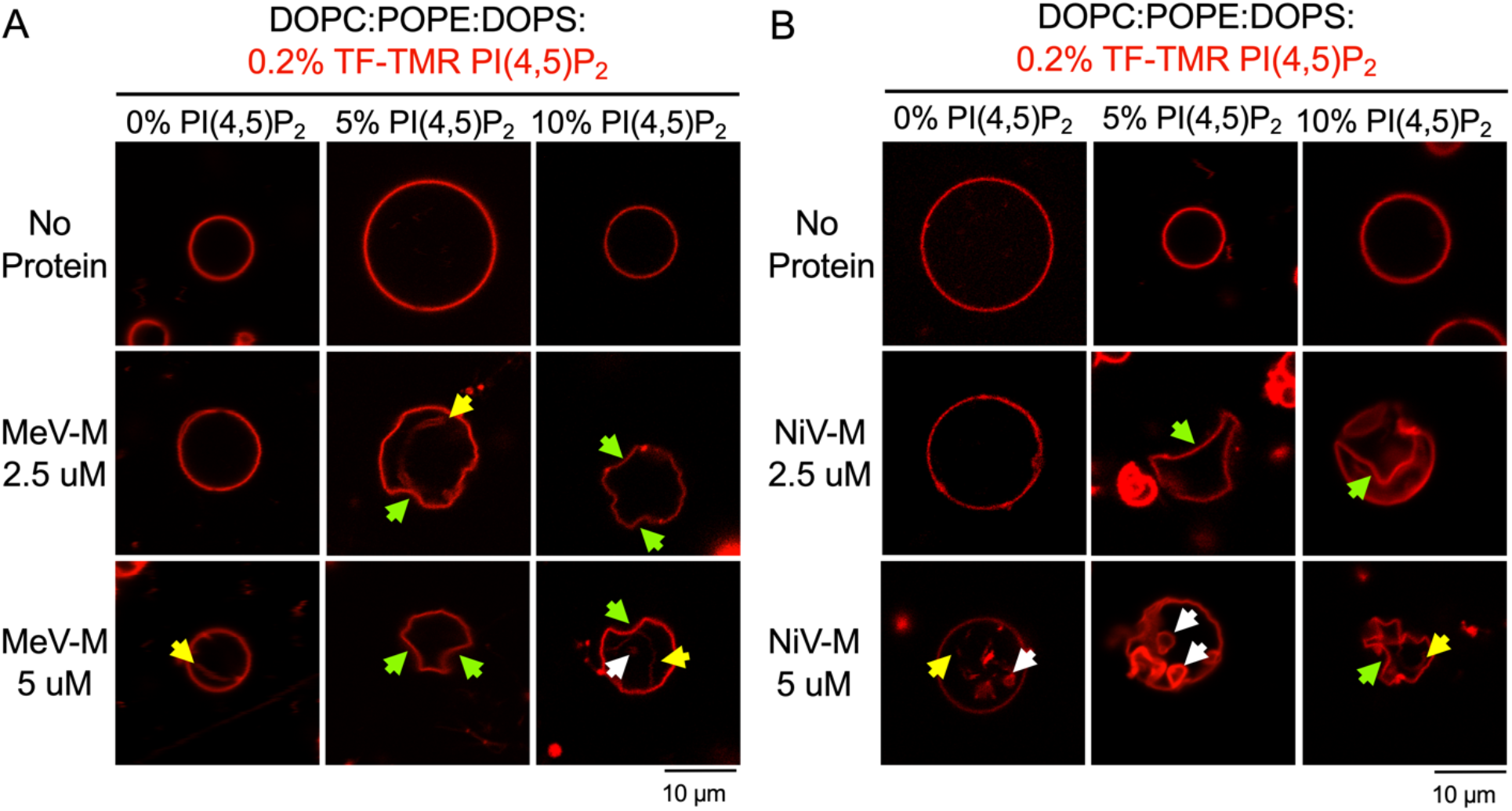
Matrix proteins induce membrane deformation in the presence of PI(4,5)P_2_. (A & B) Representative CLSM images of GUVs comprising DOPC, POPE, DOPS and 0 (left), 5 (middle), or 10 (right) mol% PI(4,5)P_2_ with TF-TMR PI(4,5)P_2_ (0.2 mol%; red fluorescence) after incubation with no protein (top), 2.5 µM (middle), or 5 µM (bottom) MeV-M (A) or NiV-M (B). No membrane deformation occurred without MeV-M or NiV-M. Filamentous (yellow arrow), spherical (white arrow), or flat (green) protrusions into the GUVs interior appeared after adding MeV-M or NiV-M. Scale bars=10 µm. CLSM: confocal laser scanning microscopy.

### Interaction with PI(4,5)P_2_ induces matrix protein lattice assembly

We next sought to determine if interaction with PS and PI(4,5)P_2_ would induce M protein lattice polymerization. nsEM of vesicles composed of POPC, POPS, and PI(4,5)P_2_ incubated with MeV- or NiV-M illustrated formation of long, ordered filaments (Figure 5A & B). These filaments formed only in the presence of PI(4,5)P_2_, suggesting that higher-order oligomeric assembly requires this lipid (Figure S7A-D). Filaments comprised a helical lattice of M protein dimers with ∼12-14 M dimers per turn and a ∼27° helical twist for MeV-M (Figure 5A), and 14-16 dimers per helical turn and a ∼7.7° twist for NiV-M (Figure 5B). Projecting 2D class averages onto 3D cylinders and docking X-ray structures into the 3D volumes showed that MeV- and NiV-M polymerization involves interactions between adjacent dimers and those in rows immediately above and below (Figure 5C & D). The spacing and arrangement of M dimer subunits in these filaments is consistent with M protein assemblies in tomographic reconstructions of the M layers of authentic paramyxoviruses (Battisti et al., 2012; Ke et al., 2018a) (Figure S7E).

**Figure 5.**
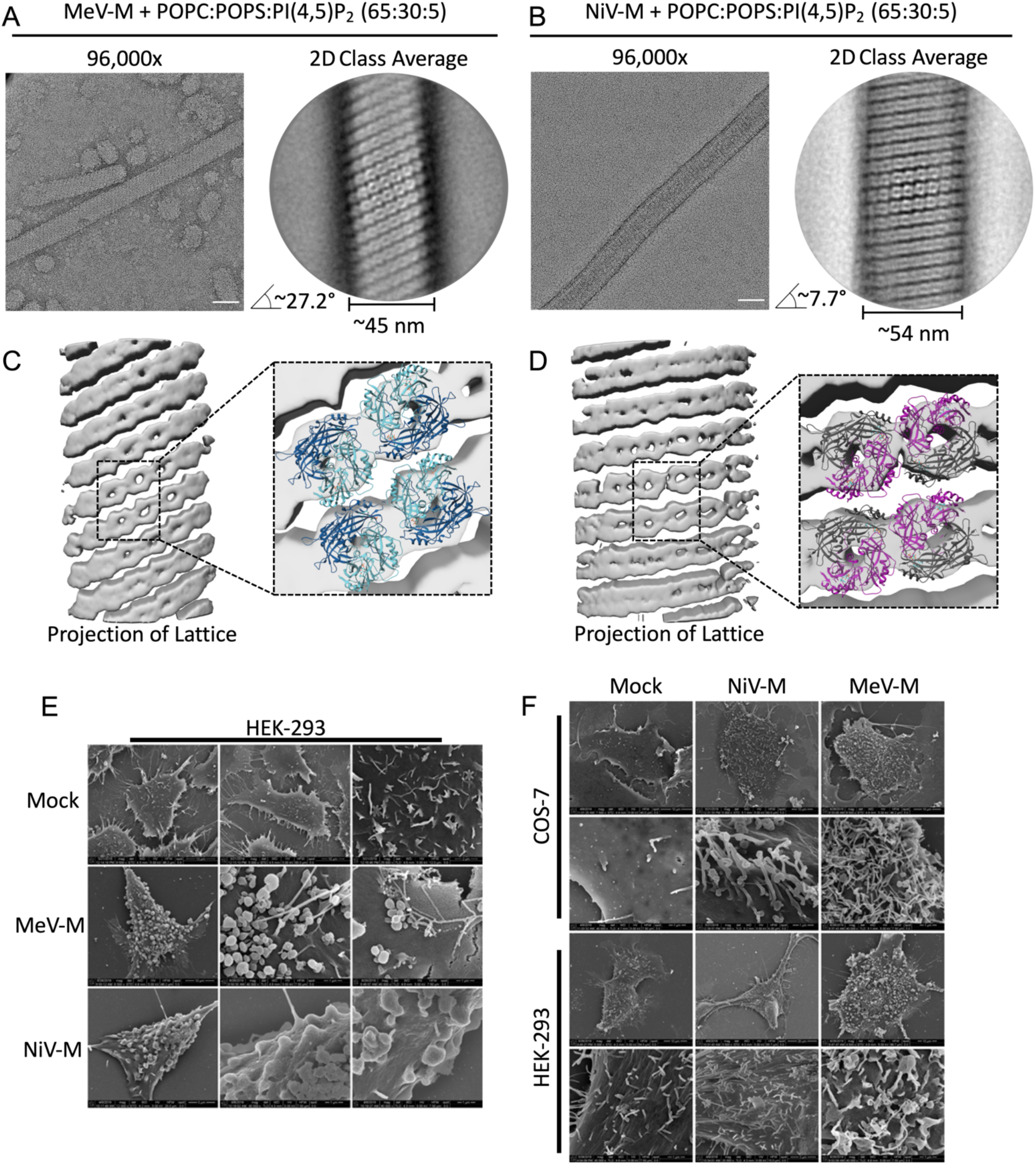
Matrix proteins self-assemble in the presence of PI(4,5)P_2_ and form spherical and filamentous extensions from cells. (A & B) *Left:* representative electron micrographs (96,000x magnification) of negatively stained liposomes comprising POPC:POPS:PI(4,5)P_2_ (65:30:5 mol% respectively) with 10 µM MeV-M (A) or NiV-M (B). Scale bars=50 nm. *Right:* representative 2D class averages of helical filaments showing twist and diameter of the helical assembly. (C & D) Isosurface representation of 2D class averages in A and B projected onto a 3D cylinder and refined without helical parameters. *Inset:* close-up of the volume (gray transparent surface) shows packing of fitted lipid-bound homology model of MeV-M or crystal structure of NiV-M bound to PI(4,5)P_2_. (E & F) SEM of COS-7 and HEK293 cells transfected with MeV- or NiV-M. (F) Representative micrographs of HEK293 cells expressing the indicated protein (or mock transfected) showing spherical particles budding from the cell surface. (G) Representative micrographs of COS-7 cells and HEK293 cells expressing the indicated protein (or mock transfected) showing filamentous particles budding from the cell surface. SEM: scanning electron microscopy.

We next used scanning electron microscopy (SEM) to delineate whether M-driven filament formation is relevant in cells and indicative of viral budding. The cell surfaces of HEK293 and COS-7 cells transiently expressing MeV- or NiV-M also exhibit abundant spherical and filamentous protrusions that are not present on the surface of mock-transfected cells (Figure 5E & F).

### Pharmacological reduction of PI(4,5)P_2_ inhibits live Measles and Nipah virus infection

The PIP5kα inhibitor ISA-2011B reduced MeV- and NiV-M localization at the PM (Figure 1H & I). To address whether inhibition of PI(4,5)P_2_ production (Figure 6A) also affects MeV and NiV in the context of authentic virus, we conducted ISA-2011B inhibitor studies in live MeV and NiV infections. ISA-2011B dose-dependently reduces replication of recombinant measles virus modified to express EGFP (rMV^KS^EGFP(3)) relative to control treated cells, with little effect on cell viability (Figure 6B & C). In NiV-infected Vero76 cells, ISA-2011B significantly reduced NiV plaque size and number (Figure 6D), suggesting that inhibitor-mediated reductions in PI(4,5)P_2_ levels are a promising strategy to reduce paramyxoviral spread.

**Figure 6.**
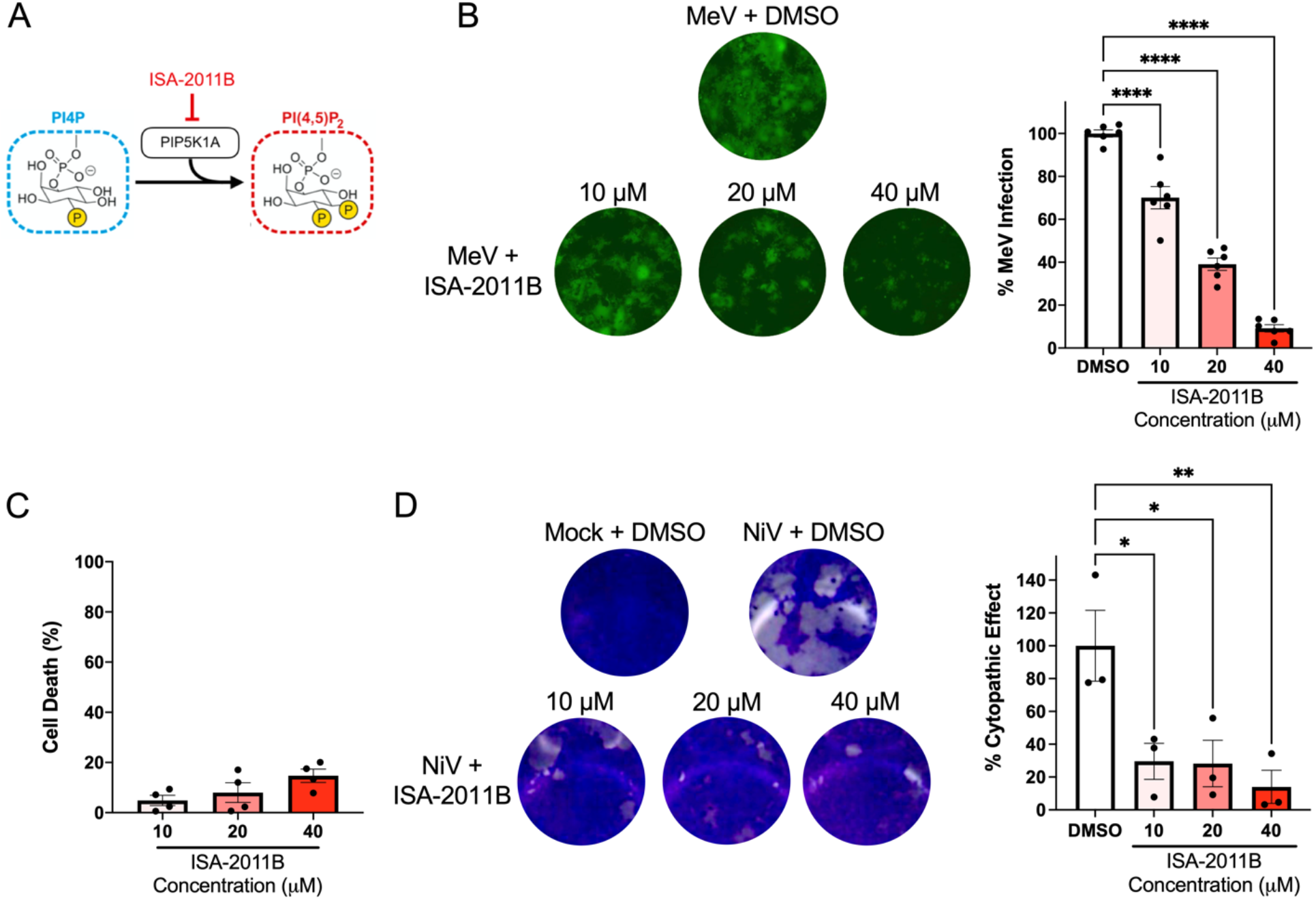
Pharmacological reduction of PI(4,5)P_2_ inhibits live measles and Nipah virus infection in cell culture. (A) Phosphatidylinositol-4-Phosphate 5-Kinase Type 1 Alpha (PIP5K1A) catalyzes formation of PI(4,5)P_2_ from PI4P and can be inhibited by ISA-2011B. (B) *Left:* representative images of Vero-hCD150 cells 42 hours after infection with rMV^KS^EGFP(3) (MOI, 0.1) and indicated ISA-2011B concentration. *Right:* Percent rMV^KS^EGFP(3) infection in ISA-2011B-treated cells normalized to DMSO-treated cells (EGFP signal as a surrogate for infection) (n=2 independent experiments performed in triplicate). (C) Vero-hCD150 cells with indicated ISA-2011B concentration for 48 hr. The y-axis denotes cell death (normalized to DMSO-treated cells) 48 hours post-treatment. Values represent mean ± SEM (n=3 independent experiments performed in duplicate). (D) *Left*: representative images of fixed and Giemsa-stained Vero76 cells 44 hours after infection with NiV (MOI 0.001) and treated with ISA-2011B as indicated. *Right:* Percentage cells exhibiting cytopathic effect after ISA-2011B treatment normalized to DMSO-treated cells. ns P > 0.05, *P ≤ 0.05, **P ≤ 0.01, ***P ≤ 0.001, ****P ≤ 0.0001 by one-way analysis of variance (ANOVA).

## Discussion

M proteins play a fundamental role in paramyxovirus assembly and release by binding cellular membranes, self-assembling, and organizing viral components at budding sites (Harrison et al., 2010). Here, we identify a critical lipid head group bound by M proteins during virus assembly, solve structures of M proteins alone and in complex with this lipid, and demonstrate dramatic conformational changes upon lipid binding that lead to membrane interactions, viral assembly, curvature, and budding. The assembly identified resembles that observed in authentic virions. We further demonstrate using mutagenesis, that dimerization, lipid coordination, and conformational rearrangement of the C-terminus are all essential for viral budding.

Taken together, these data provide the framework for a model of viral budding (Figure 7) in which monomeric M initially assembles into a dimeric confirmation (Figure 7; Step 1) present in both MeV- and NiV-M Apo crystal structures (Figure S3A & B). The large, conserved, hydrophobic dimeric interface of the Apo form gives rise to a doughnut-shaped molecule with a positively charged concave surface. Mutations in the dimer interface impede dimer formation and M-driven budding (Figure S4), supporting a role for M dimers as the basic unit for paramyxovirus matrix assembly (Battisti et al., 2012; Bringolf et al., 2017; Liu et al., 2018).

**Figure 7.**
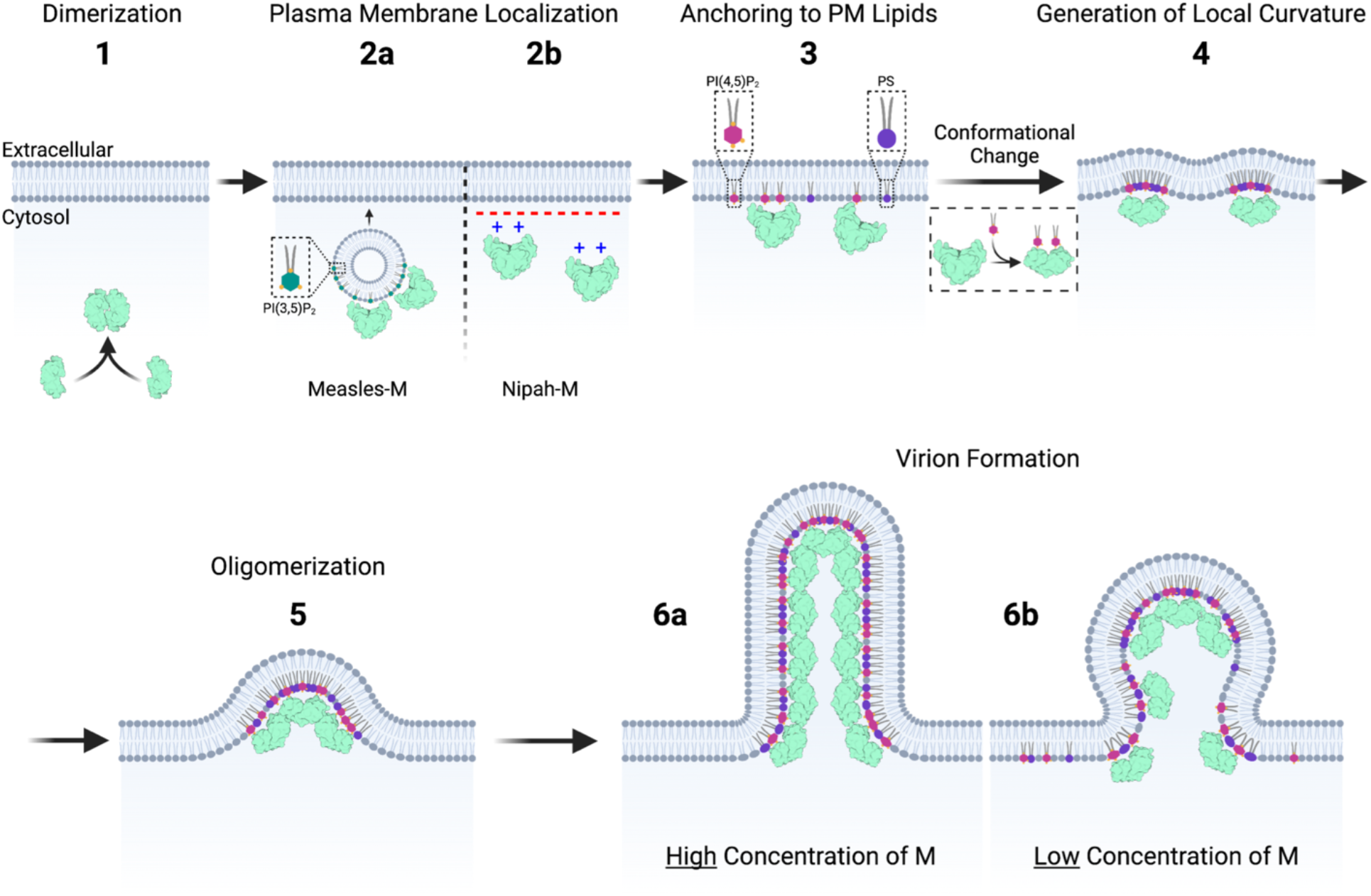
Model for membrane association and assembly of paramyxovirus M proteins. Monomeric M proteins initially assemble into a dimeric confirmation (1). M dimers then localize to the PM inner leaflet through trafficking on PI(3,5)P_2_-containing vesicles (2a) or non-specific electrostatic interactions (2b). Anchoring to the PM inner leaflet is achieved through specific binding to PI(4,5)P_2_ (magenta; 3). Interaction with PI(4,5)P_2_ causes a conformational change in M (dashed box) to open a basic patch in the NTD that then interacts with other anionic PM lipids such as PS (purple lipid). The M surface transitions from concave to flat to drive spontaneous local membrane curvature (Step 4). Membrane deformation caused by one M protein creates strong, long-range attraction between membrane-bound M proteins promoting large-scale oligomerization and matrix lattice formation that further potentiates membrane deformation (5). M clustering and oligomerization induces local asymmetric clustering of bound lipids, reducing energy needed for membrane curvature. We hypothesize that at sufficiently high surface densities, M proteins form a scaffold that shapes the membrane into a stable tubule or filament structure (6a), but at lower surface densities M proteins shape membranes into spherical structures (6b). Schematic created with BioRender.com.

Next, M proteins localize and assemble at the PM (Figure 7; Step 2). Non-specific electrostatic interactions between positively charged residues in the CTD basic patch of paramyxovirus and closely related pneumovirus M proteins were thought to drive interactions with negatively charged membranes (Battisti et al., 2012; Leyrat et al., 2014; Liu et al., 2018; Money et al., 2009). Like other paramyxovirus M proteins, we show that the CTD surface of both MeV- and NiV-M has a highly basic patch of residues (Figure S3C & D). Our assays with sphingosine indicated that MeV-M localization appears to be independent of the PM inner leaflet negative charge (Figure 1A), whereas NiV-M may rely on non-specific electrostatic interactions to reach the PM (Figure 1B). Liposome sedimentation assays indicated MeV-M interacts with PI(3,5)P_2_ (Figure 1D; S1D), which is more abundant in intracellular vesicles within the endocytic trafficking pathway than the PM (van Meer et al., 2008). Thus, MeV-M could be trafficked to the PM via association with PI(3,5)P_2_ on intracellular vesicles. The potential involvement of PI(3,5)P_2_-mediated trafficking is supported by inhibition of MeV-M PM localization by apilimod, which reduces PI(3,5)P_2_ levels (Figure S2G). In this mechanism, MeV-M already localized to the PM prior to treatment would be unaffected by apilimod, but no new MeV-M could reach the PM.

Upon arriving at the PM, M proteins must bind and anchor to the inner leaflet (Figure 7; Step 3). Although paramyxovirus M proteins have intrinsic membrane affinity (Battisti et al., 2012; Ke et al., 2018a; Riedl et al., 2002; Ringel et al., 2019; Wang et al., 2010), the detailed mechanism of M-membrane interactions is unclear. Our data indicate that both MeV- and NiV-M associate with membranes containing PS (Figure 1C; S1C), which was previously shown to be important for PM anchoring of other viral M proteins (Adu-Gyamfi et al., 2015; Bobone et al., 2017; Doktorova et al., 2017). We also show that, similar to filovirus and retrovirus M proteins (VP40 and gag, respectively) (Johnson et al., 2016; Ono et al., 2004; Wijesinghe and Stahelin, 2015), MeV- and NiV-M proteins specifically associate with membranes containing PI(4,5)P_2_ *in vitro* and in cells (Figure 1D & F-I). Importantly, the presence of both PS and PI(4,5)P_2_ induces synergistic binding by MeV- and NiV-M (Figure 1E), indicating that binding to one membrane component increases the affinity of M for the other, and for the membrane in general.

Our crystal structure of NiV-M bound to PI(4,5)P_2_ demonstrates sweeping conformational rearrangements in the M dimer (Figure 2), including expansion of the dimer interface and a transition from a concave to a flat membrane binding surface. This transition is accompanied by opening of a pocket in the CTD basic patch that receives PI(4,5)P_2_ (Figure 2A). Mutating residues that coordinate interactions with the PI(4,5)P_2_ headgroup abrogates VLP budding, supporting a function of PI(4,5)P_2_ as a direct membrane anchor and the finding that membrane association of paramyxovirus M proteins is not dependent simply on electrostatics (Figure 2F & G). Other paramyxovirus M proteins also encode positively charged residues in homologous sites; mutations to these residues in canine distemper virus, a close relative of MeV, inhibit M-induced VLP formation (Kadzioch et al., 2020), suggesting that this site is structurally similar across paramyxovirus M proteins. Similarly, the importance of the C-terminus in paramyxovirus M driven budding has also been recently highlighted for PIV (Ueda et al., 2021). In addition to coordination of the PI(4,5)P_2_ headgroup, the C8-PI(4,5)P_2_ 1′ and 2′ acyl chains are both buried within a hydrophobic pocket of NiV-M. Membrane anchoring of proteins via acyl chain binding is seen for other peripheral membrane proteins (Kinnunen, 1996; Kinnunen et al., 1994; Rytömaa and Kinnunen, 1995; Tuominen et al., 2002), including the viral matrix domain of HIV-1 gag (Saad et al., 2006). Further work is therefore needed to clarify the degree to which acyl chains contribute to NiV-M-membrane anchoring.

In addition to providing a membrane anchor, we show that PI(4,5)P_2_ binding is an allosteric trigger to form a new basic patch in the NTD that could act as a secondary positively charged binding site for other PM lipids like PS, which could synergistically anchor the M protein to the inner leaflet. Consistent with this, our MD simulations with PM models (Figure 3) show that PI(4,5)P_2_-induced conformational changes increase the surface area and number of residues interacting with the PM. The PI(4,5)P_2_-bound conformation makes more hydrogen bonds with the PM and has more contacts with additional lipids, such as PS, compared to the apo-conformation (Figure 3). Several of these additional contacts involve residues in the newly-formed NTD basic patch, which is consistent with our finding that cooperative binding to PI(4,5)P_2_ and PS enhances M-membrane interaction (Figure 1E). The conformational change in the lipid-bound form could both further stabilize M proteins on the membrane and provide an allosteric mechanism to ensure PM localization, as the secondary binding site is revealed only after binding to PI(4,5)P_2_, which is primarily located in the lower leaflet of the PM (Behnia and Munro, 2005).

Formation of new virions from the PM of infected cells requires outward curvature of the inner PM leaflet (Figure 7; Step 4). MeV- and NiV-M alone deform lipid membranes to generate spherical or filamentous protrusions with negative curvature and PI(4,5)P_2_ augmented this deformation (Figure 4). Intrinsic membrane deformation activity of HIV-1 (Mücksch et al., 2017, 2019; Ono et al., 2004) and Ebola virus (Gc et al., 2016; Johnson et al., 2016) M proteins involves PI(4,5)P_2_, as do cellular proteins like inverse Bin-Amphiphysin-Rvs (I-BAR) domain proteins, which drive formation of cellular filopodia via a PI(4,5)P_2_-dependent mechanism (Mattila et al., 2007; Saarikangas et al., 2009). Our structure of NiV-M bound to PI(4,5)P_2_ shows that upon PI(4,5)P_2_ binding NiV-M adopts an almost flat membrane interacting surface with a radius of curvature similar to the I-BAR protein, Pinkbar (planar intestinal- and kidney-specific BAR domain protein) (Figure 2C). Pinkbar proteins have a nearly flat membrane-binding surface and promote formation of planar membrane sheets (Carman and Dominguez, 2018; Pykäläinen et al., 2011). Similar to I-BAR proteins, paramyxovirus M proteins could drive spontaneous local curvature of the membrane upon interaction with the inner leaflet. The multiple bonds between the individual M proteins and the negatively charged bilayer creates a strong adhesive interface and the protein shape drives outward local curvature (Figure 7; Step 4).

Next, M proteins must self-assemble to form the matrix lattice (Figure 7; Step 5). In nsEM images, we observe helical assembly of M dimers only in the presence of PI(4,5)P_2_, suggesting that conformational rearrangements induced upon interaction with PI(4,5)P_2_ favor M polymerization (Figure 5A-D). MeV- and NiV-M filament diameters are similar to those found for helical assemblies of purified M from Sendai virus (SeV; 43-56 nm), a paramyxovirus that infects mice (Heggeness et al., 1982; Hewitt and Nermut, 1977), and the closely related pneumoviruses human respiratory syncytial virus (HRSV) and human metapneumovirus (HMPV), (HRSV 29 nm and HMPV 37 nm) (Leyrat et al., 2014; McPhee et al., 2011). The subunit spacing of individual M dimers within the helical lattice of our MeV- and NiV-M filaments is also similar to tomographic reconstructions of MeV and NDV matrix layers from authentic virus (Battisti et al., 2012; Ke et al., 2018a) (Figure S7E), indicating that the packing in our *in vitro*-assembled M filaments recapitulates the packing and assembly of M in the virion and is likely highly conserved across paramyxoviruses. Notably, alignment of MeV- and NiV-M filaments during 2D classification (Figure 5A & B) required application of a circular mask to first align a small subsection of the filament, suggesting that MeV- and NiV-M helical filaments comprise locally ordered patches of M dimers, similar to those observed in tomograms of the Ebola virus matrix layer (Wan et al., 2020). Membrane-bound proteins can readily diffuse and self-organize, and membrane deformation created by one protein can, in turn, recruit a second neighboring protein (Dommersnes and Fournier, 1999; Goulian et al., 1993; Netz and Pincus, 1995). Interactions between M and the lipid bilayer may generate strong, long-range attraction between bound M proteins, leading to large-scale protein oligomerization or scaffolding.

Extensive M polymerization leads to formation of viral particles extending from the surface of cells (Figure 7; Step 6). The angle of M dimer-dimer interactions was previously proposed to contribute to membrane curvature (Battisti et al., 2012; Liu et al., 2018). M protein clustering and oligomerization could promote asymmetric clustering of their bound lipids. Increased packing of PI(4,5)P_2_ would induce local lateral asymmetry in the inner leaflet of the PM, thereby altering its physical properties to reduce energy barriers to induce membrane curvature (Janmey and Kinnunen, 2006). Additionally, M protein oligomerization could mold the underlying lipid bilayer into a larger 3D structure that protrudes from the membrane. For BAR domain proteins, local protein density dictates the 3D protrusion morphology. At sufficiently high surface densities, BAR domain proteins form a scaffold that shapes the membrane into a stable tubule or filament structure (Bonazzi and Weikl, 2019; Simunovic et al., 2016; Sorre et al., 2012), but at lower densities spherical structures form (Baumgart et al., 2011; Isas et al., 2015; Shi and Baumgart, 2015). Indeed, our nsEM demonstrates M proteins interacting with PI(4,5)P_2_ self-assemble into long filaments with helical organization of M dimers. Virion morphology could be similarly dictated by the local concentration of M proteins and PI(4,5)P_2_. SEM evidence indicates MeV- and NiV-M produce both spherical and filamentous virus-like particles at the surface of cells (Figure 5E & F). Filamentous particles have been observed for HPIV-2 and NDV paramyxoviruses (Bang, 1949; Yao and Compans, 2000) and pneumoviral HRSV and HMPV (Ke et al., 2018b; Liljeroos et al., 2013). Thus, increased abundance of M-M contacts leads to filament formation, whereas reduced M crowding results in decreased M-M contacts leading to reduced membrane curvature and formation of spherical virions (Figure 7; Step 6a & 6b). Previous work with HRSV aligns with this possibility in that the surface area of the virion membrane covered by M varies significantly depending on virus particle morphology; filamentous particles had the highest coverage (86%) and spherical viruses had the lowest (24%) (Kiss et al., 2014).

The filamentous form may remain cell-associated and confer a selective advantage in cell-to-cell spread. The elongated shape may facilitate infection of neighboring cells while evading host mucociliary clearance (Campbell et al., 2014; Roberts and Compans, 1998; Seladi-Schulman et al., 2013, 2014). Indeed, viral propagation by cell-to-cell fusion has been previously described for MeV (Duprex et al., 1999; Herschke et al., 2007; Lamb and Jardetzky, 2007) and NiV (Aguilar et al., 2006; Lamp et al., 2013).

Our data additionally indicate that decreasing cellular PI(4,5)P_2_ by the PIP5Kα inhibitor ISA-2011B causes M protein mis-localization (Figure 1H & I) and effectively inhibits propagation of both MeV and NiV in cell culture (Figure 6). To the best of our knowledge, this is the first report to provide proof-of-principle that targeting the PI(4,5)P_2_ biosynthetic pathway could be a viable broad spectrum treatment for paramyxovirus infections. Targeting lipid metabolism is an effective antiviral strategy against multiple viruses (Beziau et al., 2020; Fernández-Oliva et al., 2019). Specifically, compounds that reduce cellular cholesterol can inhibit HPIVs (Bajimaya et al., 2017). Although we found minimal toxicity for ISA-2011B in cell culture (Figure 6C), others found that targeting lipid biosynthesis can have adverse events *in vivo* (Fernández-Oliva et al., 2019; Spickler et al., 2013). For this reason, structure guided development of small molecule inhibitors that target the conserved PI(4,5)P_2_ binding site on paramyxovirus M proteins will be important for development of broad-spectrum antivirals against paramyxovirus infections.

Paramyxoviruses represent a major risk to human and animal health and it is vital that efforts continue to focus on new targets within the virus replication cycle. Although future investigations are needed to determine how M proteins, lipids, and host proteins work in concert to form virions, illuminating M structures and their membrane interactions can explain how paramyxovirus M proteins direct virion formation and provides 3D templates for the design of agents to inhibit paramyxovirus assembly and egress.

## Acknowledgements

The authors gratefully acknowledge Katie Fuller and Tanwee Alkutkar for technical assistance. Data for the NiV-M structure were collected on beamline 12-2 of the Stanford Synchrotron Radiation Lightsource. Use of the Stanford Synchrotron Radiation Lightsource, SLAC National Accelerator Laboratory, is supported by the US Department of Energy (DOE), Office of Science, and Office of Basic Energy Sciences under Contract No. DE-AC02-76SF00515. The SSRL Structural Molecular Biology Program is supported by the DOE Office of Biological and Environmental Research and by the National Institutes of Health and National Institute of General Medical Sciences (*P30GM133894*). The contents of this publication are solely the responsibility of the authors and do not necessarily represent the official views of NIGMS or NIH. Data for the NiV-M+C8-PI(4,5)P_2_ structure were collected on beamlines 23-ID-B of the Advanced Photon Source, a US Department of Energy Office of Science User Facility operated for the DOE Office of Science by Argonne National Laboratory under contract no. DE-AC02-06CH11357. Data for the MeV-M structure was obtained through use of the Lilly Research Laboratories Collaborative Access Team (LRL-CAT) beamline at Sector 31 of the Advanced Photon Source provided by Eli Lilly Company, which operates the facility. The La Jolla Institute Zeiss LSM 880 was acquired through the Shared Instrumentation Grant Program (S10): NIH S10OD021831.

## Author contributions

Conceptualization: MJN, RVS, EOS; Methodology: MJN, MLH, WBK, SH, PC, AM, LJR, BL, WPD; Validation: MJN, MLH, WBK, JY, LJR, AH, SH; Formal Analysis: MJN, MLH; Investigation: MJN, MLH, WBK, JY, LJR, AH, SH, RP, SLB, ZLS, PC; Resources: BL, AM, WPD, RVS, EOS; Data Curation: MJN; Writing - Original draft: MJN, SLS; Writing - review & editing: MJN, SLS, KMH, LJR, BL, PC, AM, WPD, RVS, EOS; Visualization: MJN, MLH, WBK, PC; Supervision: PC, AM, WPD, RVS, EOS; Project Administration: MJN, RVS, EOS; Funding Acquisition: BL, PC, AM, WPD, RVS, EOS

## Supplementary Figure Titles and Legends

**Figure S1.**
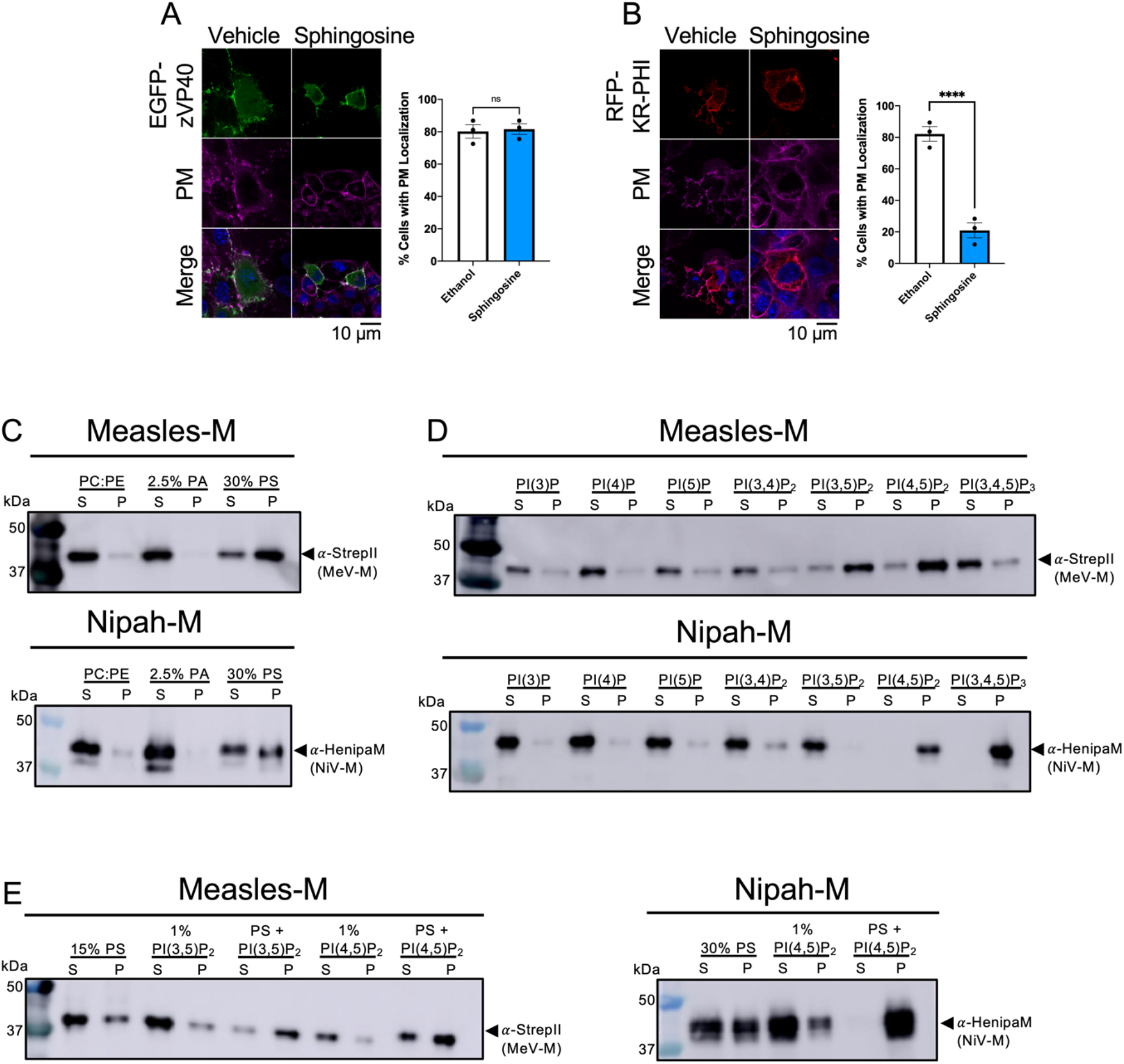
Effect of membrane charge neutralization on peripheral protein plasma membrane localization and lipid binding properties of MeV-M and NiV-M, related to Figure 1. (A & B; *Left*) Representative CLSM images of COS-7 cells expressing the indicated EGFP/mRFP fusion protein for 24 hours and subsequently treated with the membrane permeable base, sphingosine (37.5 μM or ethanol vehicle; 1:2000 v/v), for 1 hour at 37°C prior to staining with WGA AlexaFluor647 (PM stain) and Hoechst (DNA stain) and subsequent imaging on a fluorescence confocal microscope. (A) Filovirus M protein EGFP-eVP40 (B) Polycationic fluorescent probe KRφ-mRFP (A & B; *Right*) Cells with and without high fluorescence signal intensity at the PM were enumerated to determine the percentage of cells with PM localization of M. Scale bar=10 µm. At least 45 cells were counted for each replicate. Values are reported as mean ± SEM N≥135 n=3. Student’s t-test was performed to compare sphingosine treatment to the vehicle treatment group for that protein. (C-E) Liposome sedimentation assays and western blotting were performed on LUVs with varying anionic lipids and MeV- or NiV-M. Supernatant (unbound protein) and pellet (protein bound to LUVs) were separated and analyzed by western blotting using HRP-conjugated antibodies. (A) Representative western blot showing MeV-M (*Top*) and NiV-M (*Bottom*) bind to LUVs containing 30% PS but not control vesicles PC:PE or PC:PA. (B) Phosphatidylinositol binding properties of MeV- and NiV-M. Representative western blots showing MeV-M binding to LUVs containing PI(3,5)P_2_ and PI(4,5)P_2_ (*Top)* and NiV-M binding to LUVs containing PI(4,5)P_2_ and PI(3,4,5)P_3_ (*Bottom*) (C) Effect of combinations of lipids on MeV- and NiV-M affinity to liposomes. *Left*: MeV-M liposome sedimentation assays using LUVs with 15% PS +/- PI shows MeV-M binds additatively (PS+PI(3,5)P_2_) or synergistically (PS+PI(4,5)P_2_). *Right:* NiV-M liposome sedimentation assays using LUVs with 30% PS +/- PIP shows synergistic binding with LUVs containing both PS and PI(4,5)P_2_. ns P > 0.05, ***P ≤ 0.001, ****p<0.0001. CLSM: confocal laser scanning microscopy; PC: phosphatidylcholine; PE: phosphatidylethanolamine; PA: phosphatidic acid; PS: phosphatidylserine; PIP: phosphatidylinositol-phosphate; S: supernatant fraction; P: pellet fraction; LUVs: large unilamellar vesicles.

**Figure S2.**
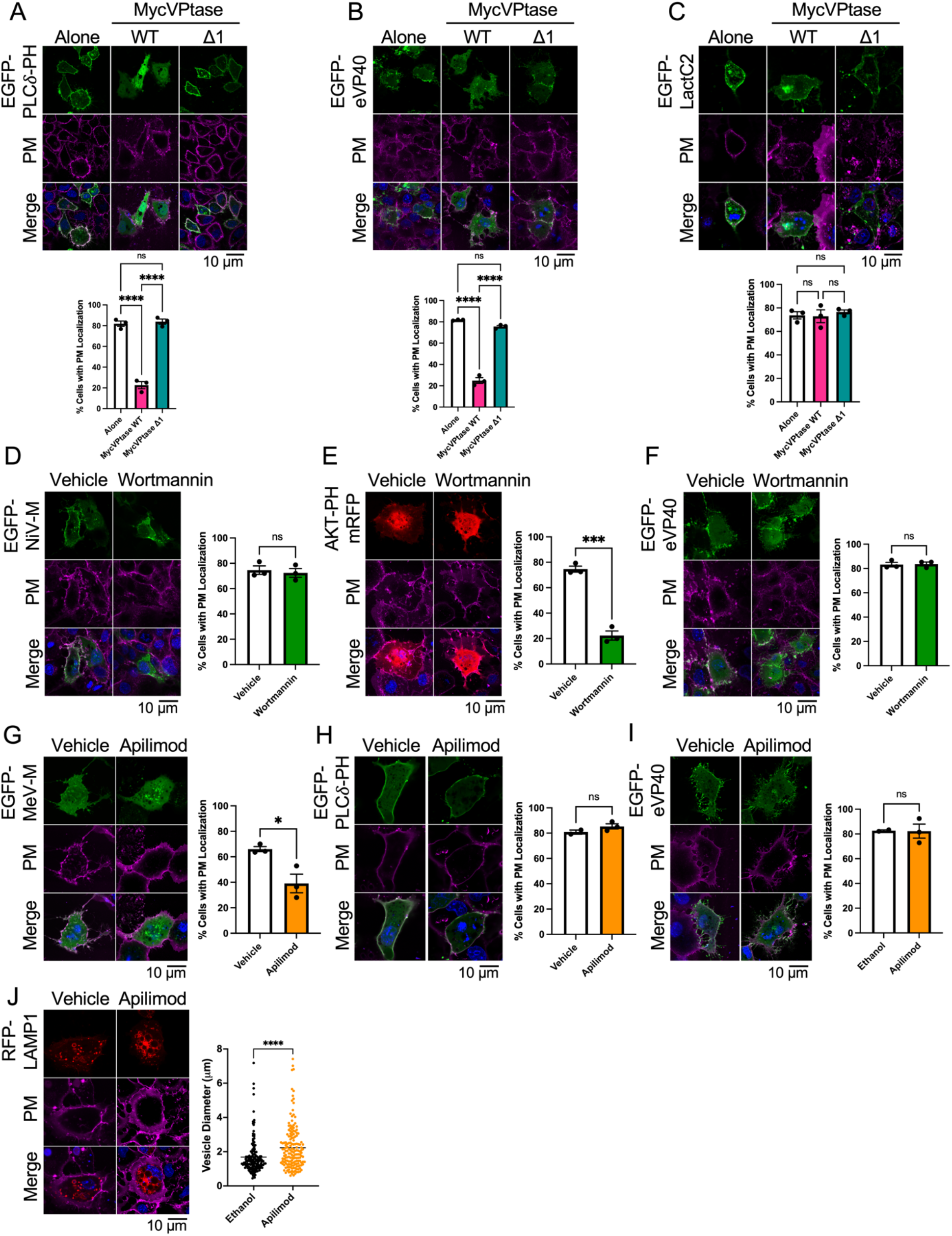
Effect of various lipid modulatory agents on peripheral protein plasma membrane localization, related to Figure 1. (A-C; *Top*) Representative confocal images of COS-7 cells expressing each indicated EGFP-fused protein alone (left panels) or with MycVPtase-WT (middle panels) or MycVPtase-Δ1 (right panels): (A) the PI(4,5)P_2_ sensor, PLCδ-PH, (B) the filoviral matrix protein, VP40 (C) the PS sensor, EGFP-LactC2. (A-C; *Bottom*) Quantification of PM localization. Cells were counted and binned based on high or no fluorescent signal at the PM and the percentage of cells with PM localization was calculated (D-F; *Left*) Representative confocal images of COS-7 cells expressing (E) EGFP-NiV-M, (F) PI(3,4,5)P_3_ sensor AKT-PH-mRFP, or (G) filovirus matrix protein EGFP-eVP40 protein treated with vehicle or wortmannin. (E-G; *Right*) Percent cells with PM localization were determined as in A-C; bottom. (G-I; *Left*) Representative confocal images of COS-7 cells expressing MeV-M (A; *Left*), PLCδ-PH-EGFP (B; *Left*), or EGFP-eVP40 (C; *Left*) +/- Apilimod treatment (G-I; *Right*) Percent cells with PM localization were determined as in A-C; bottom. (J) Analysis of Apilimod treatment on intracellular vesicle size using COS-7 cells expressing LAMP-1. (*Left*) Representative confocal images of COS-7 cells expressing LAMP-1-mCherry +/- Apilimod treatment. (*Right*) Quantification of vesicle size: intracellular vesicles were measured using ImageJ and plotted. Individual measurements are reported. A two-tailed t-test was performed. For all CLSM cells were stained 24 hr post-transfection with WGA AlexaFluor647 and Hoechst (PM and DNA stain, respectively) and imaged. Unless otherwise stated, Scale bar=10 µm. At least 45 cells were counted for each replicate. Values are reported as mean ± SEM N≥135 n=3. A one-way ANOVA with Tukey’s *post hoc* test for group comparisons was performed. ns P > 0.05, ***P ≤ 0.001, ****p<0.0001. WGA: wheat germ agglutinin; CLSM: confocal laser scanning microscopy; PM: plasma membrane; PIKfyve: phosphatidylinositol-3-phosphate 5-kinase; LAMP1: lysosomal associated membrane protein-1. PI3K: phosphatidylinositol 3-kinase; MycVPtase: Myc-5–phosphatase.

**Figure S3.**
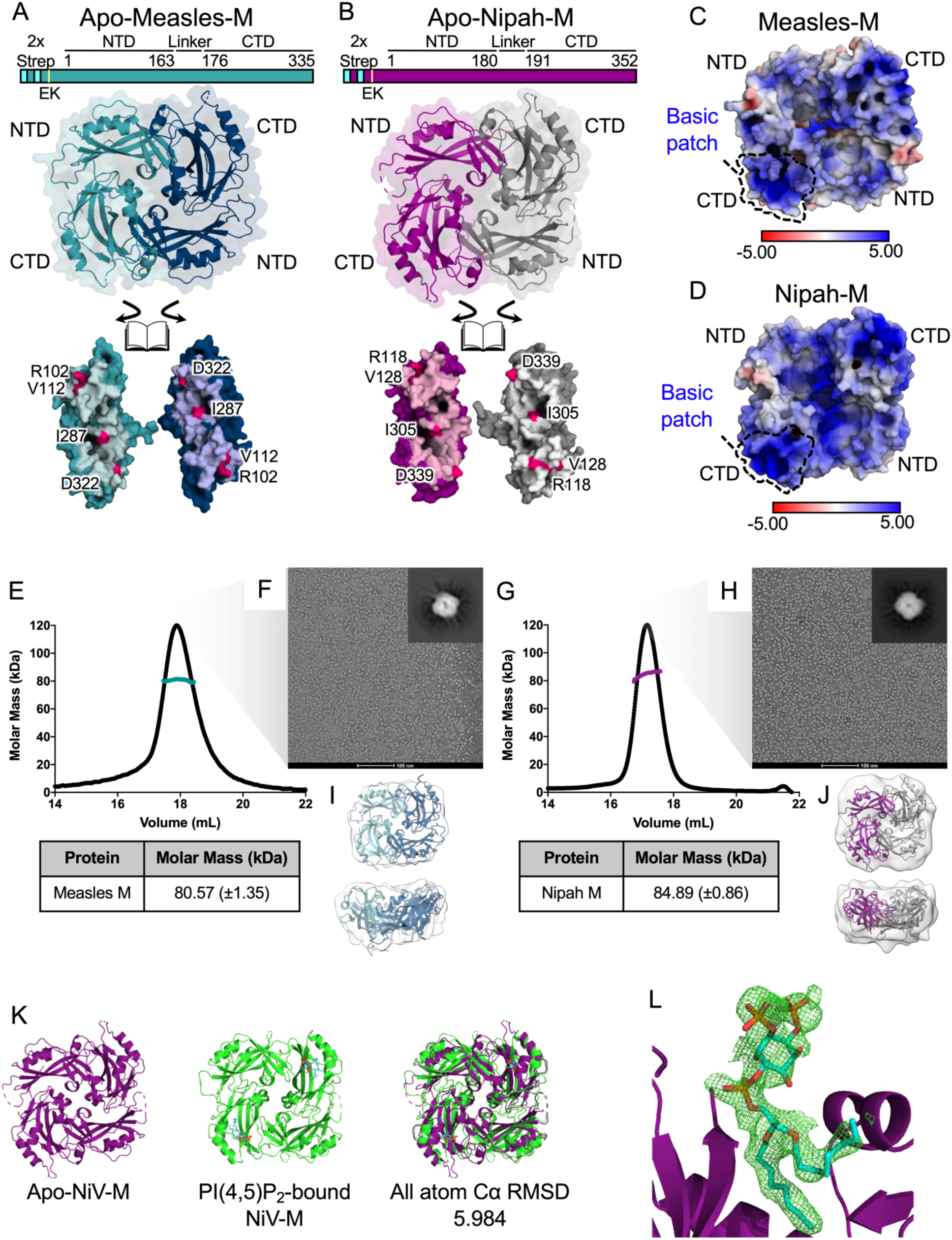
Structure and oligomerization of Apo MeV and NiV matrix proteins, related to Figure 2. (A & B) *Top:* Schematic of MeV- and NiV-M expression constructs. *Middle:* Crystal structure of M protein dimers; *Bottom:* MeV- and NiV-M dimeric interface with residues targeted for mutagenesis highlighted in magenta. (C & D) MeV- and NiV-M electrostatic surface potential on a space-filling model with positively and negatively charged regions shown in blue and red, respectively. The CTD basic patch is outlined. Electrostatic potential maps were generated using PDB2PQR and APBS software (Baker et al., 2001; Dolinsky et al., 2007). (E & G) Chromatograms for Superose 6 increase 30/300 size exclusion chromatography (SEC) experiments coupled with multi-angle light scattering (MALS). Black traces represent the Rayleigh ratio signal (a measure of light scattering) and the calculated molar masses across the eluting peaks are shown as colored dotted lines (MeV-M: teal; NiV-M: purple). The average molar mass values demonstrating that the proteins exist as dimers in solution are listed in the tables. (F & H) Uranyl acetate (UA)-stained electron micrographs of purified MeV-M (F) and NiV-M (H) displaying dimeric oligomers (magnification 75,000x). Inset shows a representative 2D class average of particles (I & J) Single particle reconstruction of MeV- and NiV-M from (F) and (H), respectively, produced donut-shaped structures consistent with dimers. Crystal structures of dimeric Apo MeV- and NiV-M were docked into EM densities. (K) Structural alignment of Apo-NiV-M (purple) and C8-PI(4,5)P_2_-bound NiV-M (green) shows significant changes to the overall architecture. The dimer structures were aligned as “dimers” using all atoms at the Cα in the respective PDB files to obtain the alignment and the indicated r.m.s.d values. (L) Fo–Fc electron density map (in green mesh) using data from C8-PI(4,5)P_2_-bound NiV-M crystals and phases calculated by omitting the C8-PI(4,5)P_2_ from the refined model. The map is contoured at 3 sigma. The refined C8-PI(4,5)P_2_ (in stick) model is superimposed onto the omit map for comparison.

**Figure S4.**
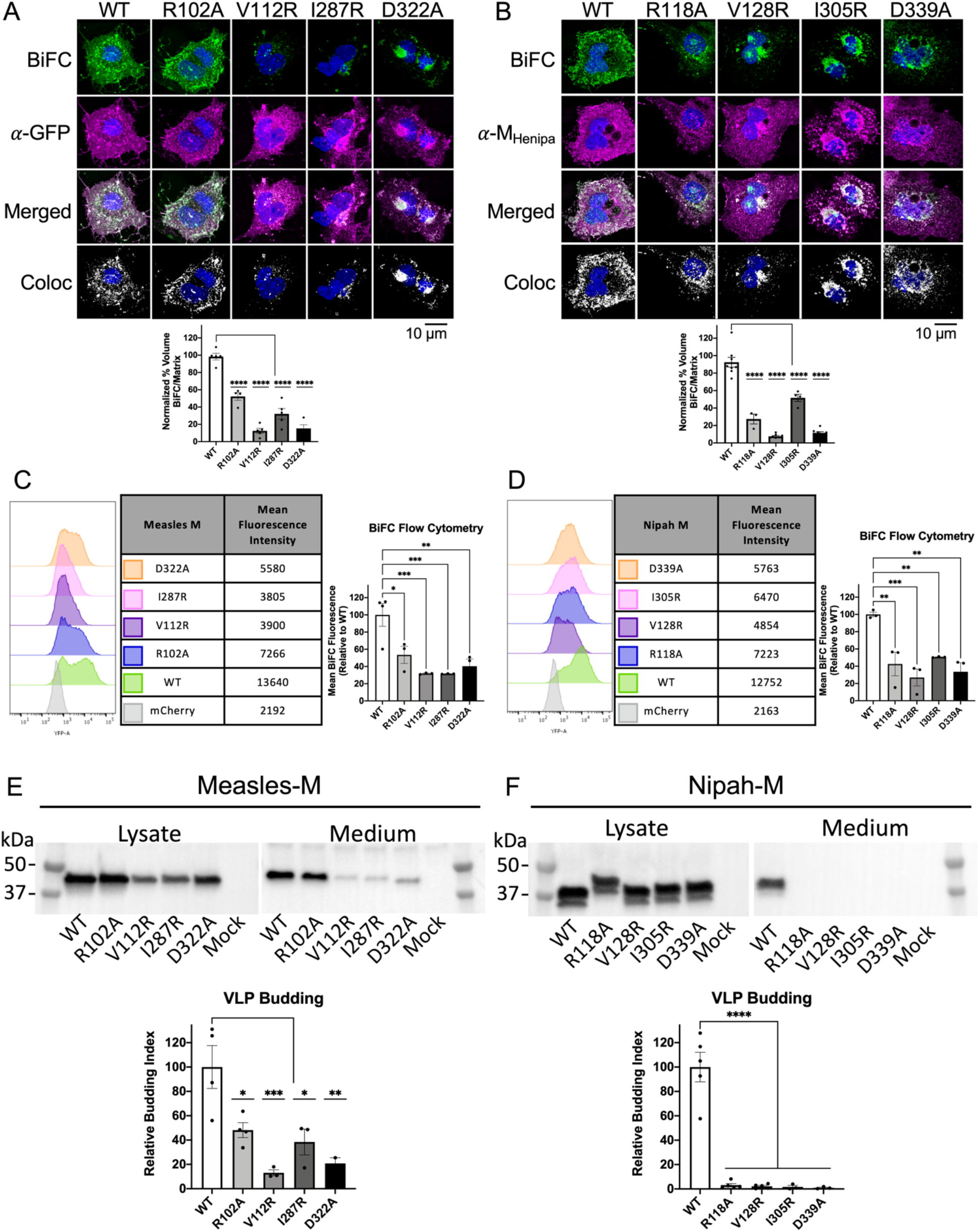
Mutations disrupting dimerization of MeV-M or NiV-M prevent virus-like particle formation, related to Figure 2. (A & B; *Top*) Representative maximum intensity projection of 3D confocal micrographs of COS-7 cells transfected with MeV-M (*Left*) or NiV-M (*Right*) wildtype or mutant BiFC matrix constructs. Oligomerization of M fused to split N- and C-terminal fragments of the fluorescent Venus protein irreversibly reconstitutes a functional fluorophore. At 24 hours post-transfection, cells were fixed and counterstained with Hoechst to visualize nuclear DNA (Blue); polyclonal anti-EGFP or anti-NiV-M antibodies were used to visualize MeV-M (G) and NiV-M (H) respectively (magenta). BiFC fluorescence is pseudo-colored green. 3D colocalization of BiFC signal with total M signal is shown in white. Scale bar=10 µm. (A & B; *Bottom*) Total 3D BiFC fluorescence/cell normalized to the total matrix fluorescence in that cell. Normalized BiFC/Matrix is plotted for each cell expressing WT or mutant M proteins (n≥3) (C & D; *Left*) Representative BiFC flow cytometry experiments for WT or mutant MeV-M (C) or NiV-M (D). Data are shown as histograms of log fluorescence intensity and colored according to the M construct. Non-aggregated mCherry+ live cells were gated and the Venus (YFP) mean fluorescence intensity (MFI) was calculated from over 10 thousand cells. Representative MFI are shown in the table on the right. (C & D; *Right*) Flow cytometry of BiFC fluorescence of WT or mutant with MFI for mutants normalized to that of WT (N≥10,000 cells/experiment). Data is from three independent experiments. (E & F; *Top*) Representative Anti-Flag (E; MeV-M) or anti-NiV-M (F; NiV-M) western blots of cell lysate and purified virus-like particles from COS-7 cells 24 hours post transfection with the indicated WT and mutant M protein constructs. (E & F; *Bottom*) VLP budding of WT and mutant M proteins with normalized budding index calculated from immunoblot intensities Unless otherwise stated, values are reported as mean ± SEM of three independent experiments. *P ≤ 0.05, **P ≤ 0.01, ***P ≤ 0.001, ****P ≤ 0.0001 by ANOVA with Tukey’s *post hoc* test for group comparisons. BiFC: bimolecular fluorescence complementation; MFI: mean fluorescence intensity; VLP: virus-like particle; WGA: wheat germ agglutinin; PM: plasma membrane; ANOVA: one-way analysis of variance.

**Figure S5.**
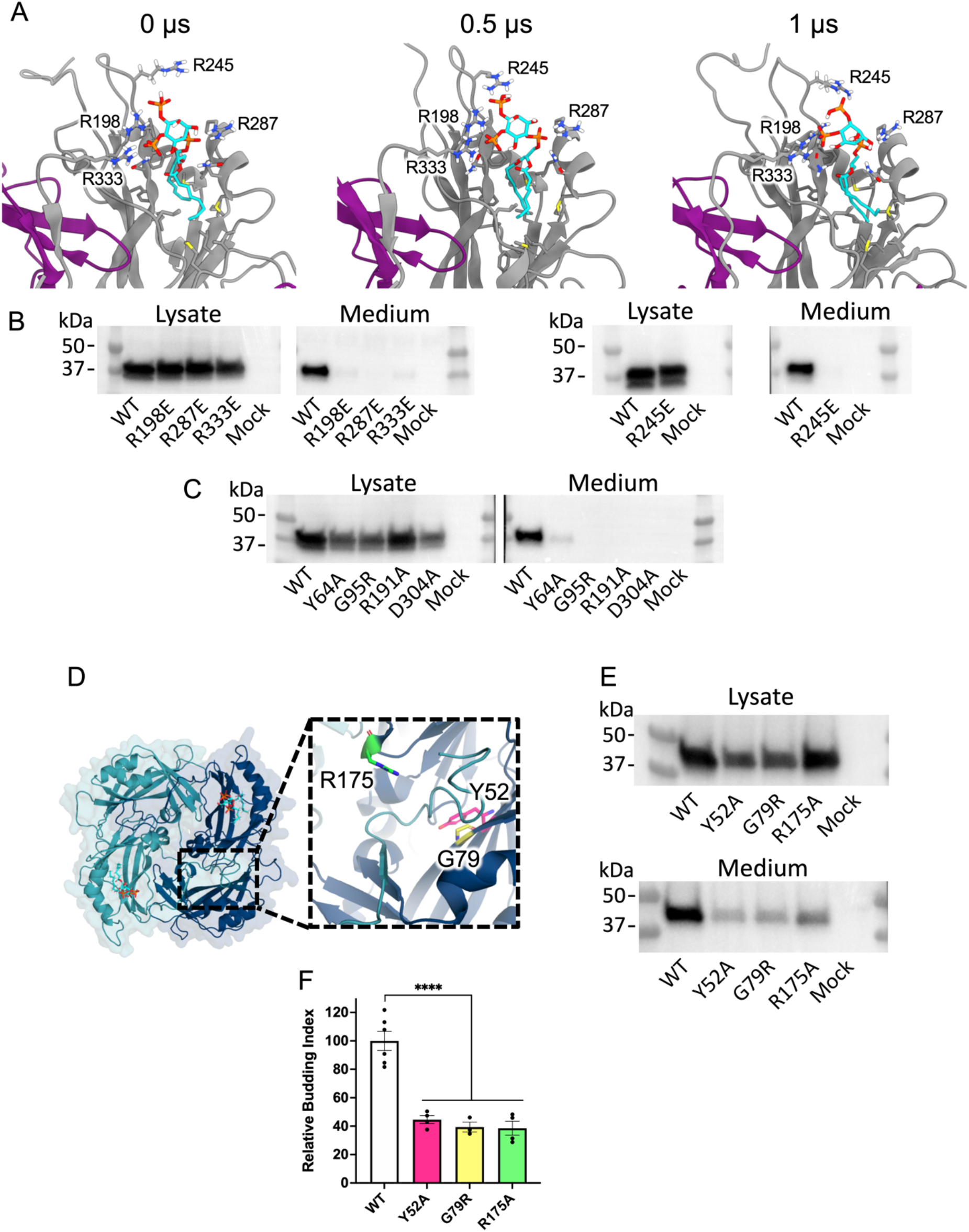
Mutations to residues involved in coordination of PI(4,5)P_2_ and the conformational change in NiV-M and predicted conformational change in MeV-M inhibit virus-like particle production, related to Figure 2. (A) Snapshots from molecular dynamics simulations of NiV-M association with C8-PI(4,5)P_2_ (Left: 0 µs, Middle: 0.5 µs, Right: 1 µs). Residues interacting with C8-PI(4,5)P_2_ are shown as sticks. By 0.5 µs R245 is interacting with C8-PI(4,5)P_2_. (B & C) Representative Anti-NiV-M western blots of cell lysate and purified virus-like particles from COS-7 cells 24 hours post transfection with the indicated WT and mutant M protein constructs. (D) Homology model of C8-PI(4,5)P_2_ bound MeV-M. Protomers are colored teal and cerulean and C8-PI(4,5)P_2_ molecules are colored cyan, orange, and red (for carbon, phosphorus, and oxygen atoms). Right inset shows a close-up view of the conformational change in the C-terminal residues (teal) extending across the hydrophobic pocket in the NTD of the adjacent protomer (cerulean). Coordinating residues in the hydrophobic pocket are highlighted according to their color in (F) and shown as sticks. (E) Representative anti-Flag (MeV-M) Western blots of cell lysate and purified VLPs from COS-7 cells 24 hours post-transfection with the indicated constructs. (F) Comparison of VLP budding between WT and mutants targeting residues coordinating the C-terminal conformational change of MeV-M. Normalized budding index was calculated from immunoblot integrated intensities. Data is the summary of three independent experiments. Values represent mean ± SEM. ****P ≤ 0.0001 by one-way ANOVA.

**Figure S6.**
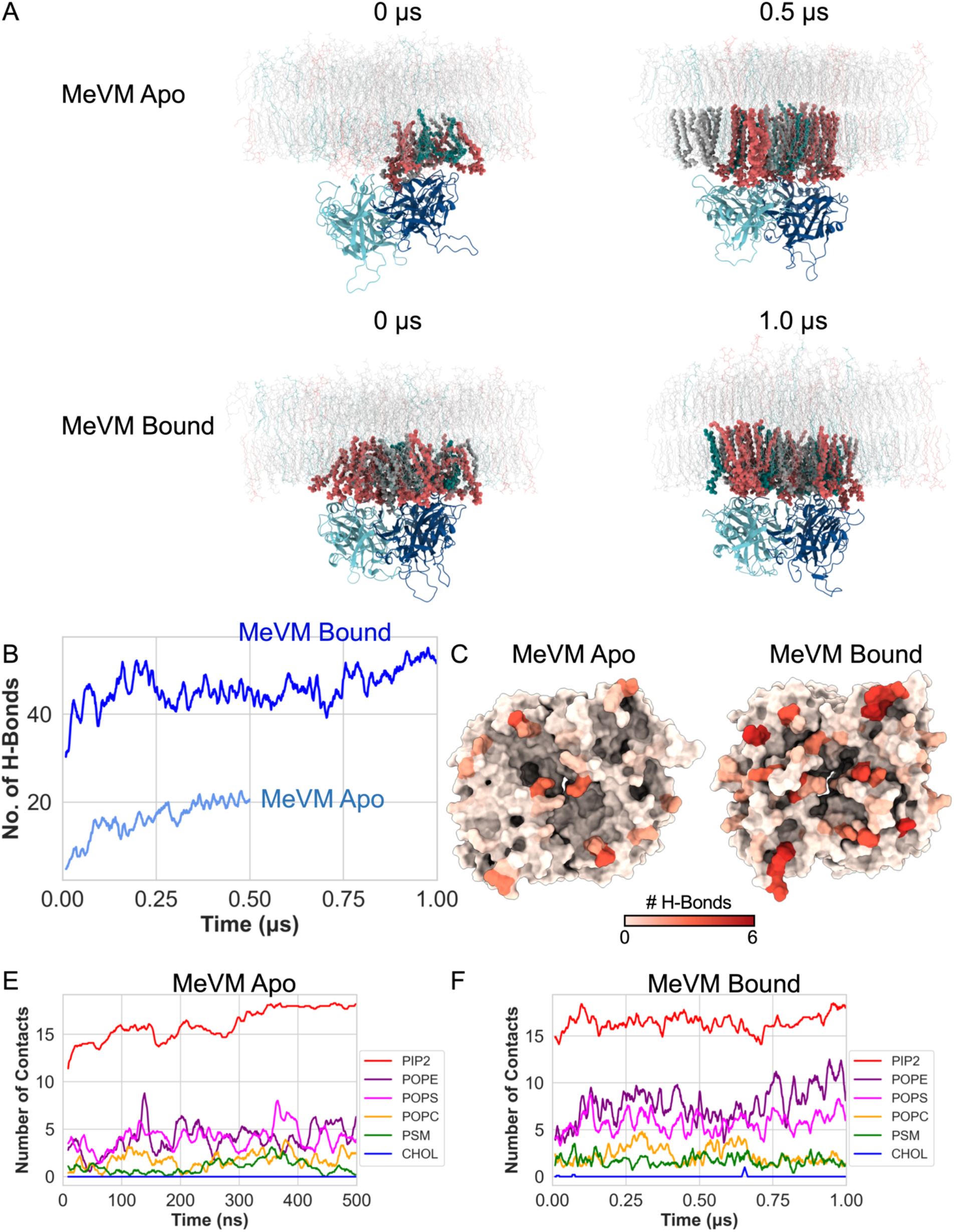
Molecular dynamics simulations of MeV-M interaction with the plasma membrane, related to Figure 3. Initial and final snapshots of MeV-M association with the plasma membrane in the (A) Apo and (B) PI(4,5)P_2_ bound conformation observed in the simulations. A homology model of the MeV-M bound conformation was generated using the bound conformation of NiV-M. The left panels display representative initial configurations and the right panels display representative final configurations. Lipids interacting with protein are highlighted (PI(4,5)P_2_ in red; PS in green; Cholesterol, POPC, POPE, and palmitoylsphingomyelin in grey). (C) Time evolution of the number of hydrogen bonds formed between membrane lipids and the Apo (light blue) or PI(4,5)P_2_-bound (dark blue) conformation of MeV-M. (D) Surface model of Apo-MeV-M (left) and PI(4,5)P_2_-bound MeV-M (right) with residues colored by the number of hydrogen bonds made with the membrane within the last 200 ns of the simulation. (E & F) Time evolution of hydrogen bonds between various lipid types and the Apo-MeV-M conformation (E) or the PI(4,5)P_2_-bound conformation (F).

**Figure S7.**
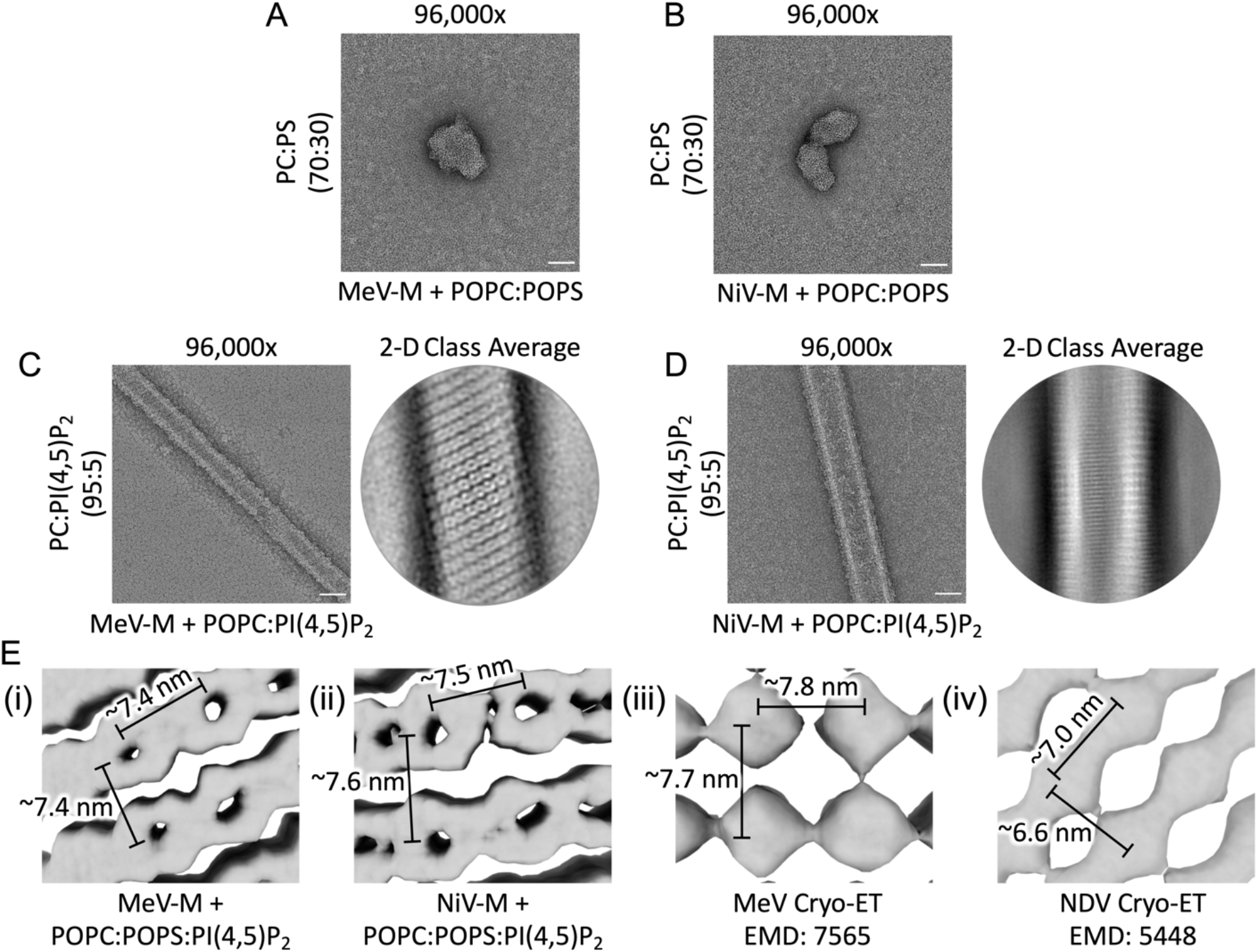
Assembly of the M protein lattice with liposomes containing PI(4,5)P_2_ is consistent with that found in virions, related to Figure 5. (A & B) Representative electron micrographs (96,000x magnification) of negatively stained liposomes composed of POPC:POPS (70:30 mol% respectively) in the presence of 10 µM MeV-M (A) or NiV-M (B). (C & D; *Left*) Representative electron micrographs (96,000x magnification) of negatively stained liposomes composed of POPC:PI(4,5)P_2_ (95:5 mol% respectively) in the presence of 10 µM MeV-M (C) or NiV-M (D). (C & D; *Right*) Representative 2D class averages of helical filaments MeV-M (C) and NiV-M (D). Scale bars=50 nm. (E) Comparison of M protein lattice from purified MeV-M incubated with POPC:POPS:PI(4,5)P_2_ liposomes (i), NiV-M incubated with POPC:POPS:PI(4,5)P_2_ liposomes (ii), Measles virus matrix lattice from Edmonston strain (EMD-7565; iii), and NDV matrix lattice from strain B1 (EMD-5448; iv). Spacing of M-protein dimers is similar across each matrix lattice.

**Movie S1. Transition of NiV-M from the Apo conformation to the PI(4,5)P_2_-bound conformation, related to** Figure 2. The rearrangement of the C-terminus is highlighted in red. The separation of beta sheet 1 and alpha helix 2 are highlighted in yellow and orange, respectively. The animation was generated in Pymol by morphing the structures between the Apo conformation and the PI(4,5)P_2_ bound conformation and back to the Apo conformation.

**Movie S2. Molecular dynamics simulations of the Apo and PI(4,5)P_2_ bound conformation of NiV-M interacting with the plasma membrane, related to** Figure 3. *Top left*: side view of the Apo NiV-M conformation. *Bottom left*: top view of the Apo NiV-M confirmation with membrane omitted for visualization. Simulation of the Apo confirmation with the membrane is 500 ns. *Top right*: side view of the PI(4,5)P_2_-bound conformation of NiV-M. *Bottom right*: top view of the PI(4,5)P_2_-bound conformation of NiV-M with membrane omitted for visualization. Simulation of the PI(4,5)P_2_-bound conformation with the membrane is 1000 ns. Positively charged residues interacting with membrane lipids are shown in blue, negatively charged in red, polar in green, and hydrophobic in white. Membrane lipids contacting protein residues are shown in khaki with stick representation.

**Movie S3. Molecular dynamics simulations of the Apo and modelled PI(4,5)P_2_ bound conformation of MeV-M interacting with the plasma membrane, related to** Figure S6. *Top left*: side view of the Apo MeV-M conformation. *Bottom left*: top view of the Apo MeV-M confirmation with membrane omitted for visualization. Simulation of the Apo confirmation with the membrane is 500 ns. *Top right*: side view of the modelled PI(4,5)P_2_-bound conformation of MeV-M. *Bottom right*: top view of the modelled PI(4,5)P_2_-bound conformation of MeV-M with membrane omitted for visualization. Simulation of the PI(4,5)P_2_-bound conformation with the membrane is 1000 ns. Positively charged residues interacting with membrane lipids are shown in blue, negatively charged in red, polar in green, and hydrophobic in white. Membrane lipids contacting protein residues are shown in khaki with stick representation.

**Table S1.**
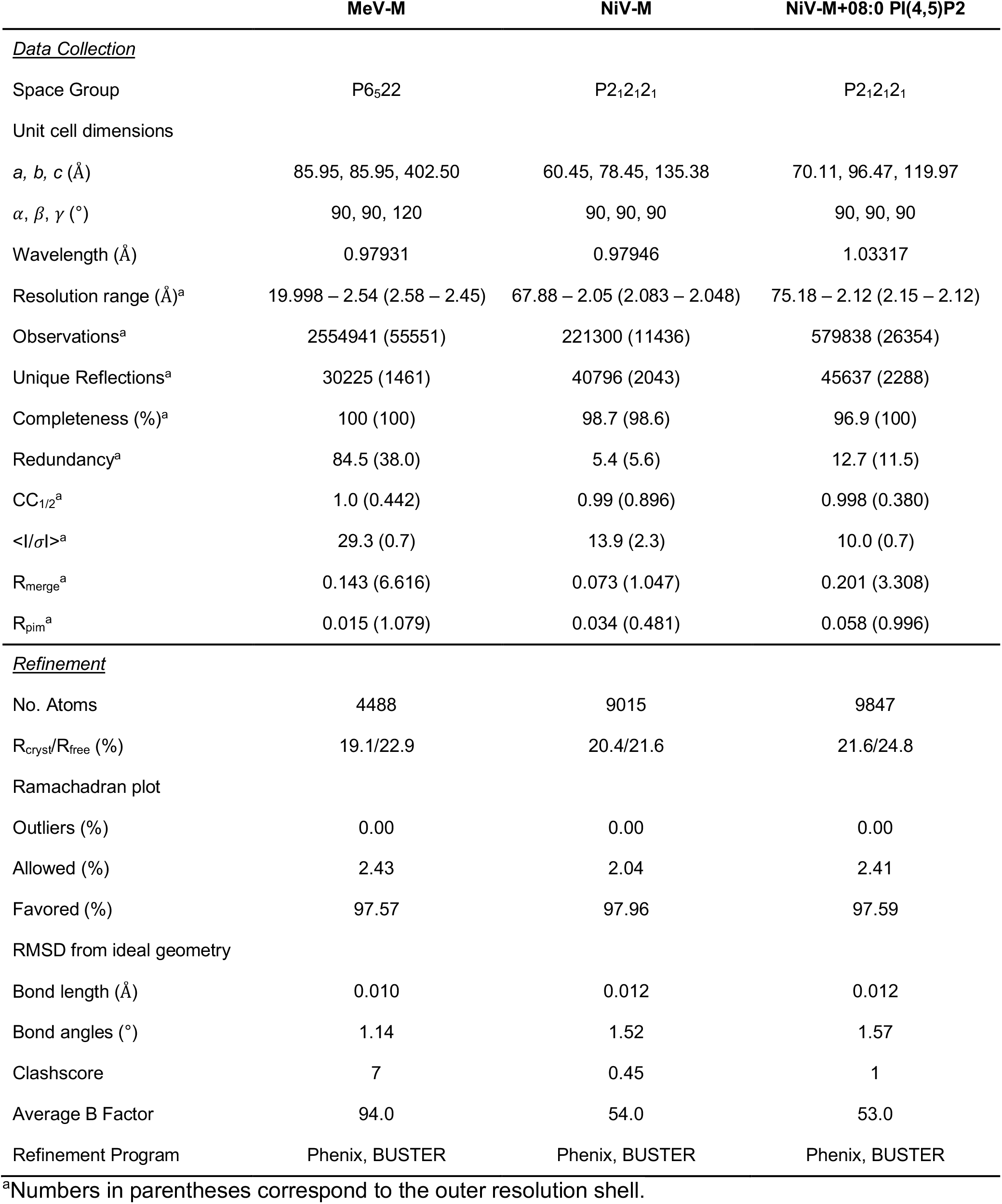
Crystallographic data collection and refinement statistics.

## Lead Contact and Materials Availability

Further information and requests for resources and reagents should be directed to and will be fulfilled by the Lead Contacts, Erica Ollmann Saphire and Robert V. Stahelin

## Methods

### Plasmids

Codon optimization and cloning of HA tagged and 3X-Flag-EGFP-tagged NiV-M is described in (Wang et al., 2010). Codon optimization and cloning of 3X-Flag-tagged and 3X-Flag-EGFP-tagged MeV-M is described in (Pentecost et al., 2015). Constructs for bimolecular fluorescence complementation (BiFC) analyses were generated with split Venus residues 1–172 (VN173) and 155– 238, A206K (VC155) (Kerppola, 2008; Shyu et al., 2006). VC155 fused to NiV-M was previously described (Pentecost et al., 2015). VN173 fused to NiV-M was generated by exchanging Ub from the previously described VN173-Ub (Pentecost et al., 2015) for NiV-M using the Gibson Assembly Cloning Kit (New England BioLabs). MeV-M BiFC constructs were made by exchanging NiV-M fused to VN173 or VC155 for MeV-M using the Gibson Assembly Cloning Kit (New England BioLabs). All mutant plasmids were generated using the Q5 site-directed mutagenesis kit using PAGE-purified mutagenesis primers designed using the online NEBase Changer primer design tool (New England BioLabs). The mCherry plasmid was created by cloning mCherry into the multiple cloning site of pTriEx-3 (Novagen) using the Gibson Assembly Cloning Kit (New England BioLabs). The EGFP-VP40 plasmid was prepared as described previously (Adu-Gyamfi et al., 2012). LactC2-EGFP and KRΦ-mRFP were kind gifts from Sergio Grinstein (University of Toronto). PLCδ-PH-EGFP was a gift from Tamas Balla (NIH). AKT-PH-mRFP was from Tobias Meyer (Stanford University). Myc-5-phosphatase-WT (MycVPtase-WT) and Myc-5-phosphatase-Δ1 (MycVPtase-Δ1) were kind gifts from Philip Majerus (Washington University). All plasmids were transformed into DH5α competent cells and maxi-preps were prepared using a Qiagen endotoxin free maxi-prep kit.

### Cell lines

HEK293 (CRL-1573, ATCC), HEK293T (CRL-3216, ATCC), COS-7 (CRL-1651, ATCC), and Vero76 (CRL-1587, ATCC) cells were cultured in Dulbecco’s modified Eagle’s medium containing L-glutamine (DMEM, Invitrogen, Carlsbad, CA) supplemented with 10% (v/v) fetal bovine serum (Omega Scientific, Tarzana, CA) and 1% (w/v) penicillin-streptomycin solution. Vero cells (CCL-81, ATCC) stably expressing the MeV receptor, human signaling lymphocyte activation molecule F1 (SLAM-F1 or CD150; Vero-hSLAM cells) were cultured in Advanced Modified Eagle’s Medium (MEM) supplemented with 10% (v/v) FBS and 2 mM GlutaMAX (Gibco-BRL). Stable expression of hSLAM was maintained using 400 g/L G418 (Sigma) (Ono et al., 2001). Cells were maintained at 37°C in a humidified atmosphere with 5% CO_2_. Insect Sf9 (*Spodoptera frugiperda*) (ThermoFisher Scientific; 11496015) cells were cultured in Sf-900 II serum free medium (ThermoFisher Scientific; 10902088) and maintained with shaking at 27°C.

### Production of viral stocks

Recombinant baculovirus was generated using the Bac-to-Bac expression system (Invitrogen) according to the manufacturer’s instructions. Briefly, the pFastBacDual vector containing an EGFP fragment under the control of the P10 promoter and MeV-M or NiV-M gene fragments under the control of the polyhedron promoter were transformed into *Escherichia coli* DH10Bac competent cells to generate bacmid DNA by site-specific transposition. Colonies bearing the recombinant bacmid were selected based on a blue/white screening method on an Luria Bertani (LB) agar plate containing kanamycin (50 µg/ml), gentamicin (7 µg/ml), tetracycline (100 µg/ml), IPTG (isopropyl-β-D-thiogalactopyranoside) (40 µg/ml), and 5-bromo-4-chloro-3-indolyl-ɑ-D-galactopyranoside (X-Gal) (100 µg/ml). The recombinant bacmid containing either the MeV-M or NiV-M gene was amplified, isolated, and transfected into Sf9 (*Spodoptera frugiperda*) cells cultured in Sf-900 II medium (Life Technologies), using PEI. The passage 1 (P1) viral stock was collected 5 days post-transfection and was used for one additional round of virus amplifications in order to produce a high-titer P2 virus stock. The recombinant measles virus, rMV^KS^EGFP(3), is based on a wild-type genotype B3 virus and its generation and propagation has been previously described (Lemon et al., 2011). The NiV_Malaysia_ strain used in this study was propagated as previously described (Moll et al., 2004).

### Lipids

All lipids were purchased from Avanti Polar Lipids, Inc. (Alabaster, AL) and used without further purification. 1-palmitoyl-2-oleoyl-glycero-3-phosphocholine (POPC; #850457), 1-palmitoyl-2-oleoyl-sn-glycero-3-phosphoethanolamine (POPE; #850757), 1,2-dioleoyl-sn-glycero-3-phosphoethanolamine-N-(5-dimethylamino-1-naphthalenesulfonyl) (dansylPE; #810330), 1-palmitoyl-2-oleoyl-sn-glycero-3-phosphate (POPA; #840857), 1-palmitoyl-2-oleoyl-sn-glycero-3-phospho-L-serine (POPS; #840034), 1,2-dioleoyl-sn-glycero-3-phospho-(1’-myo-inositol-3’-phosphate) (PI(3)P; #850150), 1,2-dioleoyl-sn-glycero-3-phospho-(1’-myo-inositol-4’-phosphate) (PI(4)P; #850151), 1,2-dioleoyl-sn-glycero-3-phospho-(1’-myo-inositol-5’-phosphate) (PI(5)P; #850152), 1,2-dioleoyl-sn-glycero-3-phospho-(1’-myo-inositol-3’,4’-bisphosphate) (PI(3,4)P_2_; #850153), 1,2-dioleoyl-sn-glycero-3-phospho-(1’-myo-inositol-3’,5’-bisphosphate) (PI(3,5)P_2_; #850154), L-α-phosphatidylinositol-4,5-bisphosphate (brain PI(4,5)P_2_; #840046) and 1,2-dioleoyl-sn-glycero-3-phospho-(1’-myo-inositol-3’,4’,5’-trisphosphate) (PI(3,4,5)P_3_; #850156) were used for liposome sedimentation assays. D-erythro-sphingosine (sphingosine; #860490) and 1-oleoyl-2-(6-((4,4-difluoro-1,3-dimethyl-5-(4-methoxyphenyl)-4-bora-3a,4a-diaza-s-indacene-2-propionyl)amino)hexanoyl)-sn-glycero-3-phosphoinositol-4,5-bisphosphate (TopFluor® TMR PI(4,5)P_2_; #810384) were used for cellular imaging experiments. 1,2-dioleoyl-sn-glycero-3-phosphocholine (DOPC; #850375) and 1,2-dioleoyl-sn-glycero-3-phospho-L-serine (DOPS; #840035) were used for giant unilamellar vesicles (GUVs). All lipid stocks were prepared in CHCl_3_ (with the exception of PIPs which were stored in CHCl_3_:Methanol 10:1) and stored at −20°C. C8-PI(4,5)P_2_ (#850185) was used for crystallization experiments.

### Liposome Preparation

Large unilamellar vesicles (LUVs) were prepared by combining POPC, POPE and dansylPE with either phospholipids (PA: 2.5%; PS: 30%) or phosphoinositides (2.5%) and dried under a steady stream of N_2_. In each experiment, addition of negatively charged lipids was compensated with an equal mol% decrease in POPC, while POPE (9%) and dansylPE (1%) were held constant. As ∼100% NiV-M was located in the pellet fraction of LUVs with 2.5% PI(4,5)P_2_ in Figure 1D, for lipid cooperation studies the amount of PI(4,5)P_2_ was reduced to 1% in order to observe an increase upon addition of PS to the LUVs. Furthermore, MeV-M binding to LUVs saturated at ∼65% of protein in the pellet fraction (Figure 1C & D), therefore the PS content was reduced to 15% (from 30%) and PI content was reduced to 1% (from 2.5%) in order to detect an increase in binding when both PS and PI species were incorporated into the LUVs. On each experimental day, lipid films were brought to RT and hydrated in extrusion buffer (250 mM Raffinose pentahydrate (Fisher Scientific; ICN10279725), 50 mM TRIS 150 mM NaCl, pH 7.4) for 45 min at 37°C. LUVs were vigorously vortexed prior to extrusion through a 200 nm Whatman polycarbonate filter (GE Healthcare). LUVs were diluted with 3x volume of LUV buffer (50 mM Tris, 150 mM NaCl, pH 7.4) and centrifuged at 50,000 x *g* for 15 min at RT. The supernatant was discarded and the pelleted LUVs were resuspended in 1x vol. of LUV buffer.

### Liposome sedimentation assays

Liposome sedimentation assays were performed as described in detail in Julkowska *et al.* 2013 (Julkowska et al., 2013). In brief, MeV- and NiV-M (0.01 mg/mL) were incubated with LUVs (400 mM) at a 1:1 vol. for 30 min at RT. Following incubation, protein-bound LUVs (pellet fraction) were separated from unbound protein (supernatant fraction) through centrifugation. Samples were then subjected to SDS-PAGE and western blotting. Equal volumes of pellet and supernatant fractions were loaded into a 10% (w/v) SDS-PAGE gel and separated at 150 V for 45 min at RT. Samples were transferred to a nitrocellulose membrane (100 V 45 min in ice) using ice cold transfer buffer. Membranes were blocked with 5% (w/v) milk-TBST (20 mM TRIS, 150 mM NaCl, 0.1% (w/v) Tween® 20, pH 7.4) and subsequently probed for their respective antibodies. Horseradish peroxidase (HRP)-conjugated antibodies were detected using Pierce enhanced chemiluminescence (ECL) reagent (ThermoFisher Scientific; PI32209) or Clarity ECL substrate (Bio-Rad; 1705060) on an Amersham imager 600 (GE Healthcare). Percent protein bound was determined using densitometry analysis in ImageJ, according to the following equation: Percent protein bound = (density pellet / density SNT+pellet)*100. Values are reported as mean ± SEM. Unless otherwise indicated, three replicates were performed in duplicate.

### Transfection

In preparation for imaging experiments, cells were washed with 1x phosphate buffered saline (PBS) and trypsinized and seeded into the appropriate vessel 24 hours prior to transfection. All transfections were performed in OptiMEM (LifeTechnologies) using Lipofectamine LTX + PLUS reagent (LifeTechnologies; 15338100) or TransIT-LT1 (Mirus) according to the manufacturer’s protocol.

### Enzymatic depletion of PI(4,5)P_2_ in Live Cells

Cells were transfected with indicated plasmids alone, or co-transfected with plasmids coding for the constitutive Myc-VPtase-WT (active), or Myc-VPtase-Δ1 (inactive) as previously described (Johnson et al., 2016; Ono et al., 2004). As a positive control, we evaluated the subcellular location of the PI(4,5)P_2_ sensor PLCδ-PH, which interacts with the PM by binding PI(4,5)P_2_. PLCδ-PH localized to the PM when expressed alone (Figure S2A; left column) or with the catalytically inactive MycVPtase-Δ1 mutant (Figure S2A; right column), but upon co-expression with wild-type MycVPtase, which reduces PM PI(4,5)P_2_ levels, PLCδ-PH localized instead to the cytosol (Figure S2A; middle column). To confirm that MycVPtase expression does not interfere with peripheral protein binding to other anionic lipids (e.g., PS) within the PM, cells expressing the PS sensor Lact-C2 alone (Figure S2C; left column) or co-expressed with either MycVPtase-WT (Figure S2C; middle panel) or MycVPtase-Δ1 (Figure S2C; right panel) were also imaged. For each condition, Lact-C2 localized to the PM in ∼75% of cells, indicating that enzymatic depletion of PI(4,5)P_2_ did not affect PS within the PM or peripheral protein binding to PS, in agreement with a previous report (Ono et al., 2004).

### Pharmacological Treatments

#### Sphingosine/Charge Neutralization Assays

Sphingosine aliquots were dried under a steady stream of N_2_ and stored at −20°C until use. On each experimental day, a fresh aliquot was thawed and resuspended in ethanol to a final concentration of 75 mM. At 24 hr post-transfection, cells were treated with either sphingosine (final concentration= 37.5 µM) or ethanol (1:2000 v/v) for 1 hr at 37°C. Cells were prepared for live cell imaging.

#### Wortmannin/PIP(3) Depletion

Wortmannin (AC328590010) was purchased from Fisher Scientific (Hampton, NH). Aliquots were prepared in ethanol and stored at −20°C until use. On each experimental day, a fresh aliquot of wortmannin (200 µM) was thawed. At 24 hr post transfection, cells were treated with either wortmannin (final concentration= 100 nM) or ethanol (1:2000 vol/vol) for 1 hour at 37°C. Cells were then prepared for live cell imaging.

#### Apilimod/PI(3)P Depletion

Apilimod (HY-14644) was purchased from MedChem Express (Monmouth Junction, NJ). Aliquots were prepared in ultra-pure grade DMSO (VWR; 97063-136) and stored at −20°C until use. On each experimental day, a fresh aliquot of apilimod (200 µM) was thawed. At 24 hr post-transfection, cells were treated with either apilimod (final concentration= 200 nM) or DMSO (1:2000 v/v) for 1-1.5 hr at 37°C. Cells were prepared for live cell imaging. As a positive control for apilimod treatment, the diameter of LAMP-1-positive intracellular vesicles was measured and found to be substantially increased in treated compared to untreated cells (2.2. µm vs. 1.7 µm), which is consistent with previous reports (Sbrissa et al., 2018) (Figure S2J).

#### ISA-2011B/PIP5Kα Inhibition

ISA-2011B (HY-16937) was purchased from MedChem Express (Monmouth Junction, NJ). Aliquots were prepared in ultra-pure grade DMSO and stored at −20°C until use. On each experimental day, a fresh aliquot of ISA-2011B (40 mM) was thawed. At 8 hr post transfection, cells were treated with either ISA-2011B (final concentration= 40 µM) or DMSO (1:1000 vol/vol) for 24 hrs at 37°C. Cells were then fixed using 4% (w/v) paraformaldehyde in PBS and stored at 4°C until imaging.

### Cellular confocal microscopy

For BiFC experiments, cells were seeded onto No. 1 glass coverslips (ThermoFisher Scientific) and the next day transfected with the indicated plasmids. At 24 hr post-transfection, cells were fixed with 4% (w/v) paraformaldehyde (in PBS) and permeabilized with 0.3% (v/v) Triton X-100. MeV-M proteins were probed using rabbit anti-EGFP polyclonal antibody (1:1000) (Invitrogen #A6455). The Venus fluorescent protein used in BiFC experiments is a genetic variant of EGFP, this polyclonal anti-EGFP also recognizes both halves of the Venus fluorescent protein (Invitrogen). NiV-M proteins were probed using rabbit anti-NiV-M antibodies (1:1000) (Wang et al., 2010). This was followed by detection with secondary anti-rabbit IgG conjugated to Alexa Fluor 647 (Invitrogen #A27040) diluted 1:1000. All antibodies were diluted in PBS+0.1% (v/v) Tween-20 (PBST). All 3D images were acquired with a Zeiss CLSM 880 Airyscan using a 63x (1.46na) objective and the 32-channel GaAsP-PMT area detector. All image stacks (on average 30 slices) were acquired with Nyquist resolution parameters using a 0.17 µm step size and optimal frame size of 1932x×1932. All 12-bit images were acquired using the full dynamic intensity range (0-4096) that was determined with the population of cells having the moderate to brightest signal expression. All Airyscan acquired images were batch processed using the Airyscan processing module utilizing ideal 3D default settings. Images were further processed for quantitative colocalization in either Zen Pro in 2D (Zeiss) and Imaris in 3D (Bitplane). Briefly, maximum intensity projections (MIPs) were generated in Zen, processed using the Zen colocalization module. Here regions of interest (ROIs) were drawn around each individual cell, previously defined minimum thresholds were input, two fluorescent channel signals were selected and the software automatically calculates pixel intensity spatial overlap coefficients between them (both manders and pearsons are scored). This method was used to quickly first assess relative patterns of association and then was followed by a more rigorous and accurate overlap assessment method in Imaris. In this software, a similar inherent colocalization module was utilized however the images were not compressed into MIPs, the analysis was based on the raw 3D signals and thus the overlap correlation coefficients scored using voxel intensity spatial overlap between compared fluorescent signals. To score for percent ratio of area and volume of oligomeric (green) *vs.* total (magenta) fluorescence, images were processed using the isosurfacing module in Imaris. Briefly, all true signals above background and autofluorescence, were auto-outlined in 3D where by all surfaced rendered clusters of BiFC positive M protein (either MeV-M or NiV-M) could be defined as ROIs with area, volume, intensity, and integrated density parameters that were exported to excel for further analysis. Here the BiFC fluorescence (pseudocolored green) per cell was normalized to the matrix fluorescence (pseudocolored magenta) in that cell. This normalized BiFC/Matrix was then plotted relative to the averaged WT BiFC/Matrix and expressed as a percentage.

For cellular lipid interaction assays, cells were seeded onto No 1.5 glass bottom 8 well plates (MatTek) at 70% confluency. Cells were post-stained with Hoechst 33342 (Fisher Scientific; PI62249) (final concentration = 16 µM) and WGA Alexa Fluor 647 (Fisher Scientific; W32466) (final concentration= 5 µg/mL) prior to imaging. Live cell imaging was performed in a live cell imaging solution (ThermoFisher Scientific; A14291DJ) or cells were fixed with 4% (w/v) paraformaldehyde (in PBS) and stored at 4°C and protected from light until imaging. Confocal imaging experiments were performed on the Zeiss LSM 880 upright microscope using a LD C-Apochromat 40x 1.1 numerical aperture water objective or Plan Apochromat 63x 1.4 numerical aperture oil objective. A 405 nm laser was used to excite Hoechst stain, and Argon lasers were used to excite EGFP (488 nm), mCherry/RFP/TopFluor® (561 nm) and WGA Alexa Fluor 647 (633 nm). Percent cells with plasma membrane localization were ratiometrically determined by the number of cells with fluorescence detected at the PM compared to the number of cells without fluorescence at the PM. In each of the three replicates performed, at least 45 cells for each condition were counted. Values are reported as the mean ± SEM.

### Flow cytometry

HEK293T cells were seeded in 6 well plates (0.8 x 10^6^ cells/well). The next day, 250 µl of transfection mix (2.5 µg total plasmid DNA and 7.5 µL TransIT-LT1) were added dropwise to each ∼80% confluent monolayer. Wells received a transfection mix containing plasmid DNA encoding WT VN-Matrix + VC-Matrix alone, WT VN-Matrix + VC-Matrix + mCherry, mutant VN-Matrix + VC-Matrix + mCherry, mCherry alone, or no DNA. Twenty four hrs post-transfection, cells were prepared for flow cytometry. Briefly, cells were washed once with ice-cold PBS, detached using 0.05% trypsin, fixed in 1% (w/v) paraformaldehyde (in PBS) and collected by centrifugation at 200 x *g* for 5 minutes. Pelleted cells were resuspended in FACS buffer (2% FBS in PBS) and stored on ice for flow cytometric analysis. Flow cytometry was performed using an LSR II Flow Cytometer (BD Biosciences) equipped with 488 and 561 laser lines. Per sample, 50,000 events were collected. FlowJo software was used to gate cells. Dead cells and cell debris were excluded from analysis using FSC *vs.* SSC, and cell aggregates were excluded by using SSC vs Pulse Width. Cells with BiFC signal or mCherry expression alone were used to determine the compensation and cut-off for cells without fluorescence. mCherry vector was co-transfected with BiFC constructs and mCherry expression was used as a positive marker to indicate successful transfection. Only mCherry+ cells were used for measurement of BiFC fluorescence. Mean fluorescence intensity (MFI) was reported as percent of positive and normalized to WT within each DNA transfection plasmid.

### VLP budding assay

Budding of virus-like particles (VLPs) into cell supernatants was detected by Western blot analyses. Wild-type/mutant MeV-M bearing a FLAG-tag and NiV-M bearing an HA-tag were cloned into pCMV and pcDNA3.1 respectively and transfected into cells using TrasnIT-LT1 transfection reagent (Mirus). VLPs were harvested 24 hours post-transfection. Cell culture medium was spun down at 3500 r.p.m. for 20 min to pellet any cells out of the media. The cleared supernatants were then ultracentrifuged at 30,000 r.p.m. with an SW-60 rotor (Beckman) for 2 hrs through a 20% (w/v) sucrose cushion-50 mM TRIS pH 7.4, 100 mM NaCl. Pelleted VLPs were resuspended in 1X NuPAGE LDS sample buffer (ThermoFisher). Cell lysates were collected by washing cells twice with PBS followed by lysis in CytoBuster (Millipore). VLPs and cell lysates were electrophoresed on SDS denaturing gels, transferred onto polyvinylidene difluoride (PVDF) Immobilon transfer membranes (Millipore), and probed with an anti-Flag or anti-NiV-M primary antibody. The relative intensities of the bands were quantified by densitometry with a ChemiDoc MP imaging system (Bio-Rad) and ImageJ. The budding index was defined as the amount of MeV-M or NiV-M in the VLPs divided by the amount in the cell lysate across each independent experiment and normalized to the average WT MeV-M or NiV-M.

### GUV Preparation and Imaging

GUVs were prepared by a gentle hydration method, as described previously (Darszon et al., 1980; Reeves and Dowben, 1969; Yamashita et al., 2002). In brief, GUVs were prepared from lipid stocks into glass round-bottom flasks by combining the appropriate volumes of DOPC, POPE, POPS, 0.2 mol% TopFluor-PC and 0.2 mol% TopFluor-TMR-PI(4,5)P_2_. In GUVs with 5 mol% PI(4,5)P_2_ and 10 mol% PI(4,5)P_2_, DOPC was reduced accordingly. GUVs were then dried gently under a steady stream of N_2_, with constant rotation of the round bottom flask. GUVs were then hydrated at 37°C overnight in hydration-buffer (250 mM sucrose, 150 mM NaCl, 20 mM HEPES pH 7.4) and used within 48 hours of preparation. Immediately prior to imaging, GUVs were diluted in glucose suspension buffer (250 mM glucose, 150 mM NaCl, 20 mM HEPES pH 7.4) at a 1:10 ratio. GUVs were placed on a 6 mm diameter chamber made from a silicon sheet using a core sampling tool (EMS #69039-60) and imaged prior to protein addition to ensure proper GUV formation and suspension. The indicated concentrations of NiV-M and MeV-M were added to the GUVs and imaging was performed at 37°C on a Nikon Eclipse Ti Confocal inverted microscope (Nikon, Japan), using a Plan Apochromat 60x 1.4 numerical aperture oil objective. The 561 nm argon laser was used to excite TopFluor-TMR PI(4,5)P_2_. In each experiment, a z-stack was taken prior to protein addition. After protein addition, a 40 min time lapse (1 frame/ 5 sec) was acquired followed by z-stacks.

### Protein expression and purification

One litre cultures of *Sf*9 cells at a density of 1.5 x 10^6^ cells per mL were infected with baculovirus at a multiplicity of infection (MOI) of 1 encoding either MeV-M fused to a twin StrepII tag or NiV-M fused to a twin StrepII tag. Forty eight hours post-infection cells were pelleted at 4,000 x *g*. Cell pellets were resuspended at 4 mL/g in buffer (MeV-M: 50 mM TRIS pH 8.0, 1 M NaCl; NiV-M: 50 mM TRIS pH 8.0, 500 mM NaCl) and lysed using a microfluidizer. The clarified lysate was bound to a 2 mL slurry of Strep-Tactin Superflow plus (Qiagen) overnight at 4°C using the batch method. The slurry was poured into a column and the flow-through buffer was collected. The resin was washed with 30 vols. of buffer, and the protein was eluted in 10-15 vols. of buffer containing 5 mM desthiobiotin. The protein was subjected to size exclusion chromatography (SEC) using a HiLoad Superdex 200 preparative grade column (Cytiva). Fractions corresponding to MeV-M or NiV-M dimer were concentrated using Amicon Ultra centrifugal filters (10,000 MWCO) and used for crystallization. NDSB-201 was added to MeV-M prior to concentration to reduce aggregation.

### Crystallization

Crystals of MeV-M were grown by the hanging drop method using a 1:1 ratio of protein to well solution with a protein concentration of 10-11 mg/mL. Well solution consisted of 19% (v/v) isopropanol, 19% (w/v) PEG 4000, 5% (v/v) glycerol, 0.095 M sodium citrate pH 5.6. Crystals formed after 2-3 days. The crystals of NiV-M were grown using the hanging drop method with a 2:1 ratio of protein (2.6 mg/mL) to well solution, and a well solution containing 0.1 M TRIS pH 7.5, 20% PEG400. Crystals grew in 2-3 weeks. To obtain crystals of NiV-M in complex with C8-PI(4,5)P_2_, purified protein (2.6 mg/mL) was mixed with 1 mM of C8-PI(4,5)P_2_ (Avanti #850185P) incubated for 1 hr at 37°C prior to setting trays. Crystals were grown using the sitting drop method using a 1:1 ratio of protein:well (0.1 M HEPES pH 7.5, 2.0 M ammonium sulfate). Initial crystals were crushed using seed beads and used in subsequent optimization trials. High quality crystals were formed using the sitting drop method with a 3:2:1 ratio of protein:well:seed in the same well solution.

### Data collection and processing

All crystals were washed in their respective well solutions prior to freezing in liquid nitrogen. Data for MeV-M were collected using the Lilly Research Laboratories Collaborative Access Team (LRL-CAT) beamline at Sector 31 of the Advanced Photon Source. Data for NiV-M were collected at the Stanford Synchrotron Radiation Lightsource (SSRL) Beamline 12-2. Data for NiV-M+C8-PI(4,5)P2 were collected at the Argonne National Laboratory Advanced Photon Source on the GM/CA 23-ID-B Beamline. Data integration and scaling were performed using the autoPROC implementation of XDS and AIMLESS (Vonrhein et al., 2011).

### Structure determination and refinement

The apo forms of MeV- and NiV-M proteins crystallize in P6_5_22 and P2_1_2_1_2_1_ space groups and diffract to 2.5 Å and 2.0 Å resolution, respectively. NiV-M bound to C8-PI(4,5)P_2_ crystalizes in the P2_1_2_1_2_1_ space group and diffracts to 2.1 Å. Isotropic data were used for model building and refinement of MeV-M, NiV-M and NiV-M+C8-PI(4,5)P_2_. Both MeV-M and NiV-M structures were determined using molecular replacement, implemented in PHENIX (Adams et al., 2010), with dimeric HeV-M (PDB: 6BK6) as the search model. The NiV-M+C8-PI(4,5)P_2_ structure was determined using molecular replacement, implemented in PHENIX (Adams et al., 2010), with the NTD and CTD of NiV-M as separate search models. Refinement of each crystal structure was done through iterative rounds of manual model building using COOT (Emsley et al., 2010), followed by refinement of the models with PHENIX and BUSTER (Adams et al., 2010; Smart et al., 2012). TLS motion was applied during refinement of all structures, with the TLS Motion Determination (TLSMD) server used to determine the TLS structure partitions (Painter and Merritt, 2006a, 2006b). Prior to refinement, 5% of each data set were set aside for the Rfree calculations (Brünger, 1992). All protein interfaces and assemblies were calculated using the PISA server at European Bioinformatics Institute (Krissinel and Henrick, 2007). Structural alignments and calculations of RMSD were carried out using the program PyMOL (http://www.pymol.org), which was used for the construction and generation of all the figures. The electrostatic surface representation was calculated in PyMOL using the APBS Electrostatics plugin (Baker et al., 2001; Dolinsky et al., 2007).

### SEC-MALS

Purified MeV-M and NiV-M were separated on a Superose 6 increase column (Cytiva) using an AKTA FPLC system. SEC was coupled in-line with the following calibrated detectors: 1) a miniDawn TREOS multi-angle light scattering (MALS) detector; and 2) an Optilab T-rEX refractive index (RI) detector. The Astra VI software was used to combine these measurements and allow the absolute molar mass of the eluting M protein to be determined (Folta-Stogniew, 2006; Wen et al., 1996).

### Scanning Electron Microscopy

For SEM experiments, cells were seeded onto collagen coated coverslips in 12-well plates at 30% confluency. Twenty four hours post transfection, cells on coverslips were fixed in 2.5% (w/v) glutaraldehyde in 0.1 M sodium cacodylate buffer, post-fixed in buffered 1% (v/v) osmium tetroxide, dehydrated with a graded series of ethanol, and dried in a Tousimis 931 critical point dryer. Dried samples were coated with platinum in a Cressington 208HR sputter coater and imaged in a FEI Nova NanoSEM 200.

### Negative-Stain Electron microscopy

MeV-M and NiV-M for single particle analysis were diluted to a concentration of 0.02 mg/mL prior to negative staining. For incubation with lipid vesicles, purified MeV-M or NiV-M (10 μM) were incubated with the indicated lipids (400 μM) for 3 days at 37°C. Three microliters of sample were each applied to freshly plasma-cleaned carbon and formvar-coated 400 mesh copper grids (Electron Microscopy Sciences). After 1 minute, excess liquid was wicked off and the grids were briefly washed on droplets of double distilled water, followed by two brief washes on droplets of a 2% (v/v) aqueous uranyl acetate solution and finally incubated on a 2% (v/v) aqueous uranyl acetate solution for 1 min. Excess stain was removed and the grids were dried thoroughly. Each sample was examined on an FEI Titan Halo 300 kV electron microscope with a Falcon 3EC camera. M proteins were imaged at 75,000x magnification and M proteins with liposomes were imaged at 96,000x magnification.

### Data processing

For single particle data sets of purified MeV-M and NiV-M, CTF correction, particle picking, 2D class averaging, and 3D reconstruction and refinement were all carried out using CryoSPARC V2.15 (Punjani et al., 2017) and the maps were displayed using ChimeraX (Pettersen et al., 2021). For single particle analysis of MeV-M and NiV-M protein:lipid filaments, CTF correction was performed using CTFFIND-4.1. Helical filaments were picked manually, segmented into ∼90% overlapping segments, and 2D classified using RELION-3.1 (He and Scheres, 2017; Zivanov et al., 2018). The initial 25 cycles of 2D classification were performed using a tight circular mask (250 Å) to improve alignment followed by an additional 10 cycles without image alignment and no circular mask. A single round of 3D refinement was done using a Gaussian noise filled cylinder (diameter calculated from the 2D class average of filament) as the starting model to project the 2D class average onto the 3D cylinder. Crystal structures were fitted into the 3D projection using ChimeraX.

### Molecular Dynamics Simulations

Membrane and protein systems for NiV-M, Apo-NiV-M, MeV-M, and Apo-MeV-M were generated with the membrane builder plugin of CHARMM-GUI (Wu et al., 2014). Modeller (Eswar et al., 2007) was used to model MeV-M using NiV-M as a template, and to fill the missing residues. The asymmetric lipid bilayer was built with POPC, POPE, POPS, PI(4,5)P_2_, PSM and CHOL in 41:7:3:3:22:20 ratios in the outer leaflet and in 10:35:15:13:4:20 ratios in the inner leaflet (Bhattarai et al., 2017; Gc et al., 2016, 2017; van Meer et al., 2008; Zachowski, 1993), with 205 total lipids in upper leaflet and 208 in the lower leaflet. Each system was neutralized by 0.15 mM NaCl and solvated with TIP3P water model. The resulting dimension of each simulation box was approximately 12 nm x 12 nm x 12 nm. The same NiV-M membrane system was used for all systems by replacing NiV-M with other proteins after structural alignment.

All-atom MD simulations were performed using the GPU version of NAMD 2.12 (Phillips et al., 2005) with the CHARMM36m forcefield (Huang et al., 2017) in explicit water solvent. The minimization and equilibration of the membrane system were performed following the 6-step protocol as given by CHARMM-GUI (Wu et al., 2014). Periodic boundary conditions were employed in all simulations. The long-range electrostatic interactions were treated using the particle mesh Ewald (PME) method (Essmann et al., 1995) and the hydrogen atoms were constrained with the SHAKE algorithm (Ryckaert et al., 1977). A Nose-Hoover Langevin-piston method was used with a piston period of 50 fs and a decay of 25 fs to control the pressure. All simulations were run at 300 K temperature with Langevin temperature coupling and a friction coefficient of 1 ps^−1^. The NiV-M and MeV-M membrane systems were run for 1 s each, whereas the Apo-NiV-M and Apo-MeV-M systems were run for 0.5 s each. For calculations of hydrogen bonds, a cut-off distance of 3.5 Å and a cut-off angle of 30° were used. Visual molecular dynamics (VMD) (Humphrey et al., 1996) was used to visualize and analyze the trajectories

### Virus inhibition assay

#### Measles virus

For MeV inhibition studies, 2 x 10^4^ Vero-hSLAM cells in 96-well plates were infected with rMV^KS^EGFP(3) at an MOI of 0.1 in the presence of ISA-2011B or DMSO at the indicated concentrations. At 42 hours post-infection, when cytopathic effect was maximal in the no inhibitor control cells, fluorescent images were acquired using a DMI3000B inverted microscope equipped with a DFC345 FX camera and LAS software (all Leica Microsystems). Capture settings were adjusted so as to have no over exposure, and identical capture settings were used for all image acquisition. EGFP fluorescence was quantified in ImageJ (Schindelin et al., 2012).

#### Nipah virus

All inhibition and infection studies with NiV were performed under biosafety level 4 (BSL-4) conditions at the Institute of Virology, Philipps University Marburg. Briefly, 2 x 10^4^ Vero76 cells seeded into 96-well plates in DMEM+5% (v/v) FCS and supplemented with the indicated concentration of ISA-2011B or DMSO were infected with NiV_Malaysia_ at an MOI of 0.001 at 37°C. At 44 hours post-infection, cells were fixed with 4% (w/v) PFA, permeabilized with ethanol and stained with Giemsa to visualize NiV spread and plaque formation as previously described (Weis et al., 2014). Plates were scanned (Triumph-Adler DCC 2930) and digitized using a Chemidoc XRS+ imaging system (Biorad). Images were converted to black and white, thresholded, segmented, and the total plaque area for each well was quantified using ImageJ software (Schindelin et al., 2012).

### Cell viability assay

To confirm that a decrease in viral infection correlated with the inhibition of viral replication and not an increase in cell death, a viability screen was run in tandem using uninfected Vero-hSLAM cells. Briefly, cells were seeded at a density of 6,000 cells/well in black, flat, clear-bottom 96-well plates (Costar) and allowed to attach overnight at 37°C. The next day, medium was replaced with 200 μl of fresh medium containing indicated concentrations of compound. Cells were cultured at 37°C for 2 days. Following incubation, cell viability was determined by the addition of 20 μl of PrestoBlue viability reagent to the culture medium. The mixture was incubated at 37°C for 1 hour and the fluorescent signal was quantified on a TECAN Spark 10M fluorescent plate reader.

### Quantification and Statistical Analysis

Statistical details of experiments, including numbers of replicates and measures of precision (standard deviation, SD) can be found in the figure legends, figures, results and methods. All analyses were performed with GraphPad Prism, version 7.

### Data and Code Availability

Structure factors and associated model coordinates of MeV-M, NiV-M, and NiV-M+08:0 PI(4,5)P_2_ have been deposited in the Protein Data Bank (http://www.rcsb.org).

